# Effector Gene Reshuffling Involves Dispensable Mini-chromosomes in the Wheat Blast Fungus

**DOI:** 10.1101/359455

**Authors:** Zhao Peng, Ely Oliveira Garcia, Guifang Lin, Ying Hu, Melinda Dalby, Pierre Migeon, Haibao Tang, Mark Farman, David Cook, Frank F. White, Barbara Valent, Sanzhen Liu

## Abstract

Newly emerged wheat blast disease is a serious threat to global wheat production. Wheat blast is caused by a distinct, exceptionally diverse lineage of the fungus causing rice blast disease. To understand genetic diversity in wheat-infecting strains, we report a near-finished reference genome of a recent field isolate generated using long read sequencing and a novel scaffolding approach with long-distance paired genomic sequences. The genome assemblage includes seven core chromosomes and sequences from a dispensable mini-chromosome that harbors effector genes normally found on the ends of core chromosomes in other strains. No mini-chromosomes were observed in an early field strain, and two mini-chromosomes from another field isolate each contain different effector homologous genes and core chromosome end sequences. The mini-chromosome is highly repetitive and is enriched in transposons occurring most frequently at core chromosome ends. Additionally, transposons in mini-chromosomes lack the characteristic signature for inactivation by repeat-induced point (RIP) mutation genome defenses. Our results, collectively, indicate that dispensable mini-chromosomes and non-dispensable core chromosomes undergo divergent evolutionary trajectories, and mini-chromosomes and core chromosome ends are coupled as a mobile, fast-evolving effector compartment in the wheat pathogen genome.

**Significance statement:** The emerging blast disease on wheat is proving even harder to control than the ancient, still-problematic rice blast disease. Potential wheat resistance identified using strains isolated soon after disease emergence are no longer effective in controlling recent aggressive field isolates from wheat in South America and South Asia. We report that recent wheat pathogens can contain one or two highly-variable conditionally-dispensable mini-chromosomes, each with an amalgamation of effector sequences that are duplicated or absent from pathogen core chromosome ends. Well-studied effectors found on different core chromosomes in rice pathogens appear side-by-side in wheat pathogen mini-chromosomes. The rice pathogen often overcomes deployed resistance genes by deleting triggering effector genes. Localization of effectors on mini-chromosomes, which are unstably transmitted during growth, would accelerate pathogen adaptation in the field.

## Introduction

Wheat blast is an explosive emerging disease capable of 100% yield losses. Little resistance is available in cultivated wheat varieties, and fungicides are not effective under disease favorable conditions (Kohli *et al.* 2011; Cruz and Valent 2017). The disease emerged in Brazil in 1985 and spread within South America, limiting wheat production **(Fig. 1)**. Wheat blast jumped continents in 2016, impacting ~15% of the total wheat area in Bangladesh on this first report (Islam *et al.* 2016; Malaker *et al.* 2016). Wheat blast has now established in South Asia, enhancing fears about further disease spread, disruption of global grain trade by this seed-borne pathogen, and endangerment of global food security. Wheat blast is caused by a wheat-adapted lineage of *Magnaporthe oryzae* (synonymous with *Pyricularia oryzae*) (Gladieux *et al.* 2018), known as the *Triticum* pathotype (MoT). MoT strains are distinct from rice pathogens in the *Oryza* pathotype (MoO) and millet pathogens in the *Eleusine* (MoE) and *Setaria* (MoS) pathotypes (**Fig. S1**). A serious turf grass disease emerged in the United States in the late 1980s, caused by the *Lolium* pathotype (MoL) with ryegrass as its major host. Although some MoL strains can infect wheat (Farman *et al.* 2017), MoT strains are distinguished as highly aggressive wheat pathogens that are so far restricted to certain countries in South America and South Asia (**Fig. 1A**).

**Fig. 1.**
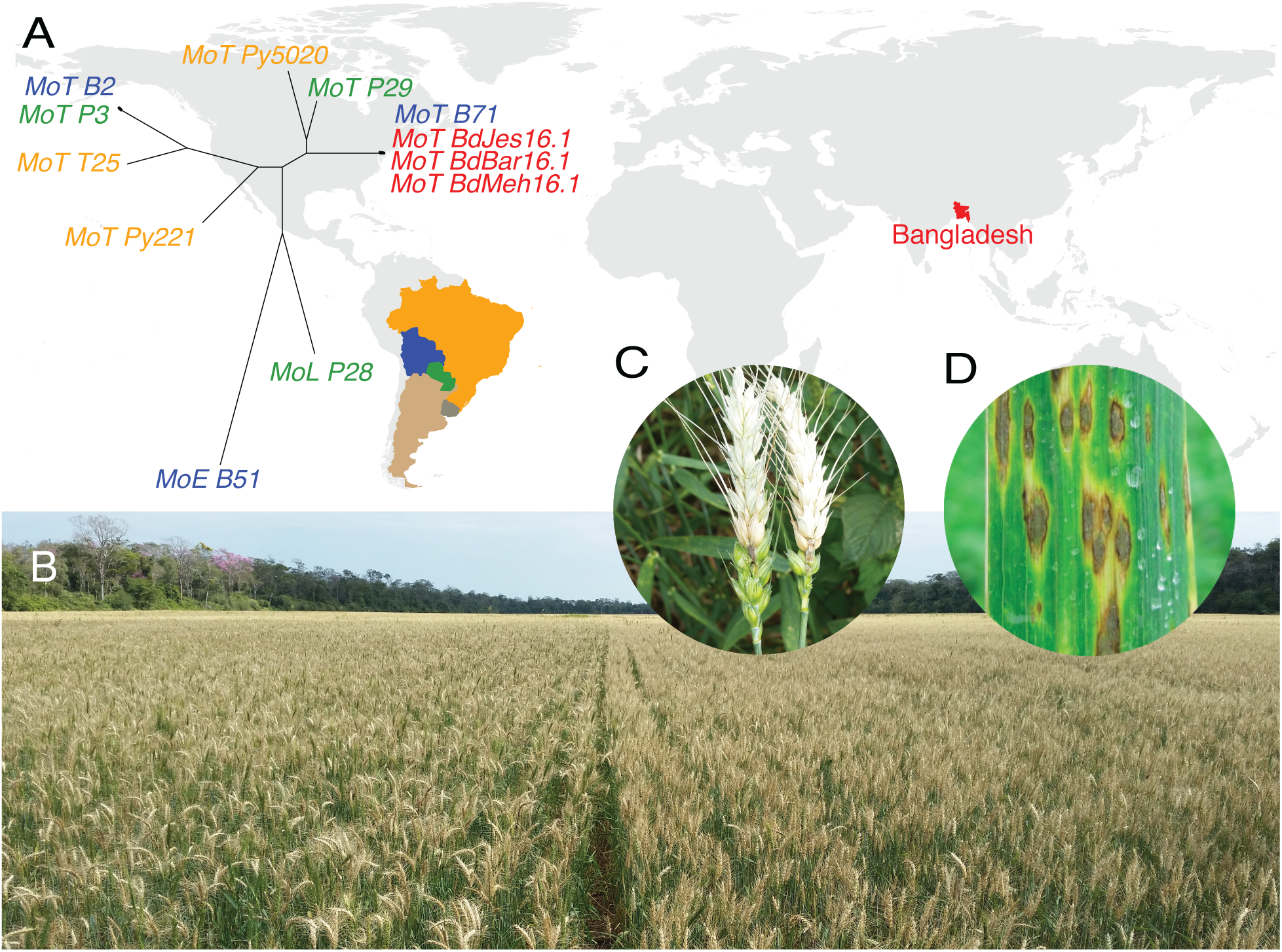
Wheat blast disease has now spread from South America and established in South Asia. (**A**) Countries with wheat blast are labeled with non-gray colors. The phylogeny contains nine strains examined in this study and three from Bangladesh. Strain names are color-coded with country colors on the map. (**B**) A wheat field in Bolivia in 2015 shows near 100% killed (straw-colored) heads. The field appeared healthy before heading. (**C**) Infected wheat heads with spikelets removed show fungus at the infection point on the stem. (**D**) Although head blast is the main symptom on wheat in the field, sporulating leaf lesions can sometimes be found on highly susceptible wheat varieties.

Although little is known about wheat blast, studies on rice blast disease have identified numerous effector genes, generally encoding small proteins that are specifically expressed *in planta* and play roles in host invasion (Giraldo and Valent 2013). Some effectors, termed avirulence (AVR) effectors, determine either rice cultivar or host species specificity through blocking infection upon recognition by corresponding cultivar- or species-specific resistance (*R*) genes and triggering hypersensitive resistance. For example, strains of several *M. oryzae* pathotypes are able to infect weeping lovegrass, *Eragrostis curvula*, unless they carry an active member of the *PWL2* gene family, host species-specific *AVR* effectors that block infection of *Eragrostis* spp. (Kang *et al.* 1995; Sweigard *et al.* 1995). Planting of wheat varieties lacking the *R* gene *Rwt3* in Brazil likely enabled MoL strains with the corresponding host species-specific *AVR* effector *PWT3* to adapt to wheat, and subsequent loss of *PWT3* function played a role in the wider emergence of the MoT subgroup (Inoue *et al.* 2017). Characterization of 12 MoO *AVR* effector genes controlling rice cultivar specificity showed most occur in transposon-rich regions of the genome, with some located near telomeres or on conditionally dispensable mini-chromosomes (supernumerary chromosomes that are unstable during vegetative growth and the sexual cycle) (Luo *et al.* 2007; Chuma *et al.* 2011; Mehrabi *et al.* 2017; Wang *et al.* 2017). Effectors are generally associated with frequent presence/absence polymorphisms between and/or within the different *M. oryzae* lineages (Chuma *et al.* 2011; Yoshida *et al.* 2016). Understanding *AVR* effector gene dynamics, including frequent deletion in response to deployed *R* genes, is key to combating the ability of the blast fungus to rapidly overcome deployed *R* genes and developing sustainable disease control.

To facilitate understanding genetic variability and evolutionary potential of the wheat blast fungus, here, a near-finished reference genome of an aggressive MoT strain was generated and compared to genomes of early and recent wheat pathogens and other host-adapted strains. We report that the genome structures of the 7 wheat blast core chromosomes have not diverged significantly from the rice blast core chromosomes. However, mini-chromosomes present in zero, one or two copies in different strains serve as a highly variable compartment for effector genes.

## Results

### A reference genome of the MoT strain B71

We sequenced and generated a near-complete genome assembly of the highly aggressive Bolivian field isolate B71 (Malaker *et al.* 2016), which exhibits high sequence similarity with MoT isolates from Bangladesh (**Fig. 1A, Table S1 and Fig. S1**). An assemblage containing 31 contigs was produced from >12.4 Gb of whole genome shotgun (WGS) PacBio long reads (**Table S2**). Genome polishing utilizing ~10 Gb Illumina sequencing data corrected 37,982 small insertions and deletions as well as 350 base-pair substitutions in the PacBio draft assembly (**Data S1**). Corrected assembled contigs were in the range of 44.2% to 52.5% GC content with the exception of a contig of 28.4%, which was predicted to be from mitochondria of B71 owing to its high similarity (99% identity) to the mitochondrial sequence of *M*. *oryzae* rice pathogen 70-15 (Dean *et al.* 2005). A circularized B71 mitochondrial sequence was obtained after removing redundant sequences at the contig ends.

We developed a novel scaffolding technology, LIEP (Long Insert End-Pair sequencing) to improve the continuity of the assembly (**Fig. 2A**). Briefly, LIEP involved construction of millions of vectors, each of which contains a unique DNA barcode pair of 22 nt and 21 nt random barcodes. Barcodes for each vector were sequenced to establish a sequence database of barcode pairs. The vectors were then used to construct clones with 20-30 kb long inserts of B71 genomic DNA flanked by the two vector barcodes. Both ends of the insert were sequenced, generating clone-end sequences with paired barcode sequences used to recover clone-end pairs. All steps were performed with pooled clones rather than individual clones. Scaffolding condensed the assembly to 12 contigs, which were then reoriented and renamed based on the MG8 genome assembly of rice pathogen 70-15 (Dean *et al.* 2005). Consequently, the final B71 genome assembly (B71Ref1) is comprised of ~44.46 Mb in seven chromosomes and five unanchored scaffolds (**Fig. 2B**).

**Fig. 2.**
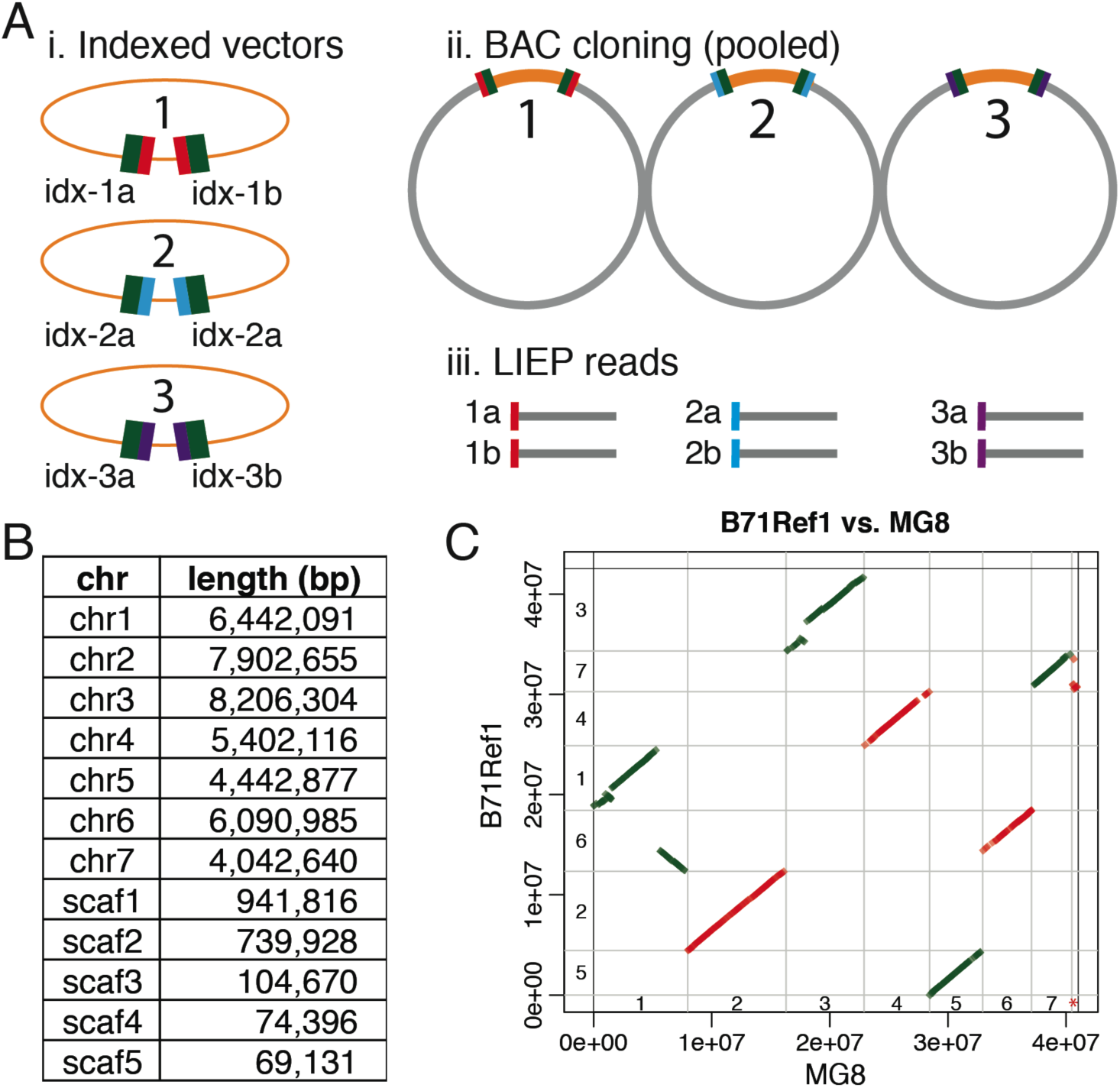
LIEP procedure and B71 assembly. (**A**) Each vector molecule contains two Illumina adaptor sequences (green) and random barcodes. The pool of barcoded vectors were pre-sequenced and are used to construct a clone library. Clone-ends, each of which included a barcode, are sequenced massively and assembled separately. Paired clone-end sequences (e.g., 1a and 1b) from the same clone are identified based on barcodes. (**B**) Lengths of 12 sequences of the B71 assembly. (**C**) A dotplot to compare the collinearity between the B71 assembly and MG8, the assembly for rice strain 70-15. Alignments with at least 10 kb match and at least 95% identity were shown. The red highlighted asterisk represents the unanchored MG8 contig (supercont8.8) that was mapped at the beginning of B71 chromosome 7.

Telomere repeat sequences (TTAGGG)_n_ or *M. oryzae* telomeric retrotransposons (MoTeRs) that integrate in telomere repeats (Starnes *et al.* 2012) were identified on both ends of chromosomes 2, 4, 5, 6, 7 and on one end of chromosome 1, indicating that B71Ref1 is a near end-to-end assembly. The B71Ref1 and MG8 assemblies show high end-to-end co-linearity for chromosomes 2, 4, 5, and 7 (**Fig. 2C**). A two-megabase rearrangement was identified between chromosomes 1 and 6, of which part of chromosome 1 of MG8 was located on chromosome 6 of B71. The rearrangement was supported by eight pairs of LIEP sequences (**Fig. S2**) and by 50 single PacBio long reads. This rearrangement is not MoT specific because it was also observed in a MoO field isolate, evidenced by a long PacBio assembled sequence spanning both chromosome 1 and chromosome 6 of MG8 (Bao *et al.* 2017). A large sequence in B71Ref1, from 1.3 to 2.9 Mb on chromosome 3, was absent in MG8. The unanchored 70-15 MG8 contig, supercont8.8, was mapped at the beginning of B71 chromosome 7, implying supercont8.8 is the missing end of chromosome 7 in the MG8 reference genome. None of five unanchored scaffolds of B71Ref1 can be mapped to MG8, with the requirement of, at minimum, a 10-kb match and 95% identity. Annotation of B71 identified 12,141 genes, with 1,726 harboring signal peptide domains (**Fig. 3**, **Data S2, S3**). Of the 248 highly conserved core set of eukaryotic genes, 243 (98.0%) orthologs from the B71 annotation were identified by CEGMA, compared to 97.6% orthologs in MG8. Therefore, completeness and annotation of the B71 genome are at least comparable to that of MG8, which was produced using Sanger sequencing and multiple technologies.

**Fig. 3.**
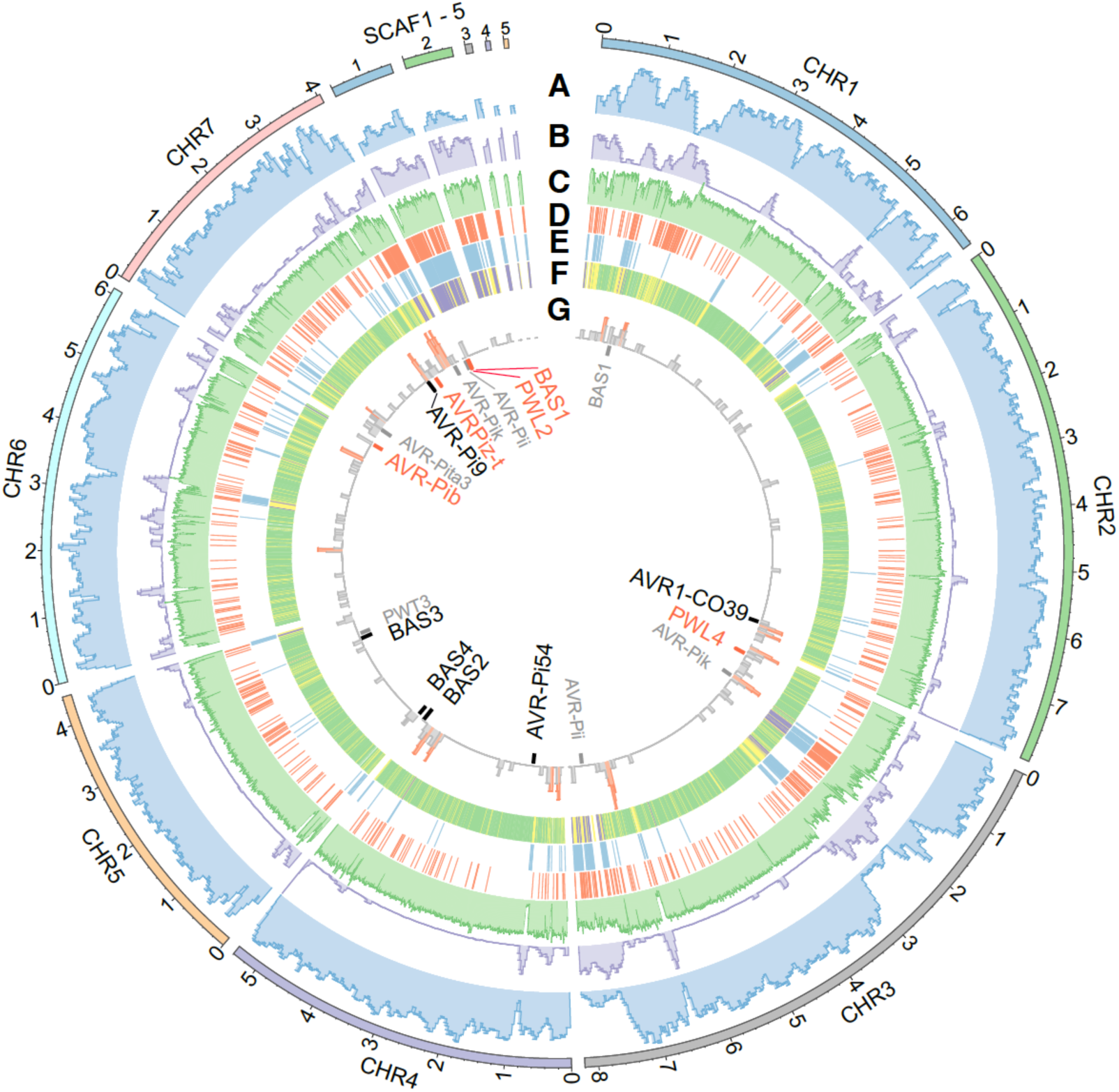
Genomic features in the B71 genome. (**A**) Density of predicted genes. (**B**) Density of repetitive sequences. (**C**) GC content. (**D,E**) Regions subjected to CNplus (**D**) or CNminus (**E**) in at least one of eight isolates (P3, B2, B51, T25, Py22.1, Py5020, P28, and P29). (**F**) CNequal regions among all isolates, CNplus or CNminus regions, and regions subjected to both changes, are colored in green, yellow, and purple, respectively. (**G**) Distribution of putative effector genes. The 100 kb regions with at least 3 putative effector genes are highlighted in red. Genes with at least 90% similarity to published effector sequences are labeled in red and black, with and without *in planta* specifical expression from our RNA-Seq analysis, respectively. Genes with a 50-90% similarity to published effector sequences are in gray.

Comparison of RNA-Seq data from MoT-infected wheat and culture-grown MoT identified 335 and 153 genes that were only expressed *in planta* and in culture, respectively (SI Materials and Methods) (**Data S4**). Secretion signal domains occurred in 173 *in planta*-specific genes, and in 18 culture-specific genes. The *in planta*-specific genes included homologs of five MoO effector genes, including *PWL2* and *PWL4* that block infection of *Eragrotis* spp. (Kang *et al.* 1995; Sweigard *et al.* 1995), *AVR-Pib* and *AVRPiz-t* that determine rice cultivar specificity (Li *et al.* 2009; Zhang *et al.* 2015), and the cytoplasmic effector *BAS1* (Mosquera *et al.* 2009) **(Fig. 3G and Data S4)**. The remaining 168 *in planta*-specific genes were considered putative effectors **(Data S5)**. Both known and putative effector genes tended to be located towards the ends of core chromosomes **(Fig. 3G)**.

### Abundant copy number variation among *M*. *oryzae* isolates

We sequenced eight additional field isolates, including less-aggressive early strain T25 from 1988 (Cruz *et al.* 2016), five other MoT strains, a MoL strain, and a MoE strain **(Fig. S1 and Table S1)** (Gladieux *et al.* 2018). A read depth approach was employed to detect genomic copy number variation (CNV) between B71 and each isolate, focusing on the identification of genomic regions with conserved copy number (CNequal), higher copy number (CNplus), or lower copy number (CNminus) in non-B71 isolates (**Fig. S3**). Among ~41.7 Mb of low repetitive regions, 36.4 Mb (87.3%) exhibited CNequal among all nine isolates. In total, 4.9 Mb (11.8%) displayed CNV between B71 and at least one other isolate, with 2.7 Mb (6.5%) being CNplus and 3.4 Mb (8.2%) CNminus (**Fig 3D, 3E, 3F**). Ten effector homologs (*PWL4*, *AVR-Pik-chr3*, *AVR-Pi54*, *BAS1-chr1, BAS2*, *BAS3*, *BAS4*, *AVR1-CO39*, *AVR-Pi9*, and *AVRPiz-t*) resided in CNequal regions (chromosome identifier added to distinguish homologs). Four (*AVR-Pii*, *AVR-Pib*, *PWL2*, and *BAS1*) were in CNminus regions and three (*AVR-Pii-scaf1, AVR-Pib*, and *AVR-Pik*) in CNplus (**Table S3**). CNV analysis of effector genes was supported by Illumina draft assemblies of the eight strains (**Table S4**). Thus, some *AVR* homologs are equal in copy number and stable across all strains, while others are subject to copy number changes. Of 1.2 Mb genomic sequences exhibiting CNplus in some isolates but CNminus in others, ~819 kb (68.5%) were from the five scaffolds (scaf1-5), which constitute only 4.3% of the genome. CNV variation of sequences in the B71 scaffolds indicated they are absent in the less aggressive MoT strain T25 (**Fig. 4A**). The P3 and B71 comparison, however, suggested that most scaffold sequences are duplicated in P3, an aggressive isolate from Paraguay in 2012 (**Fig. 4B**). In summary, extensive copy number variation was observed among *M*. *oryzae* field isolates, especially in five scaffolds that were not anchored to seven chromosomes.

**Fig. 4.**
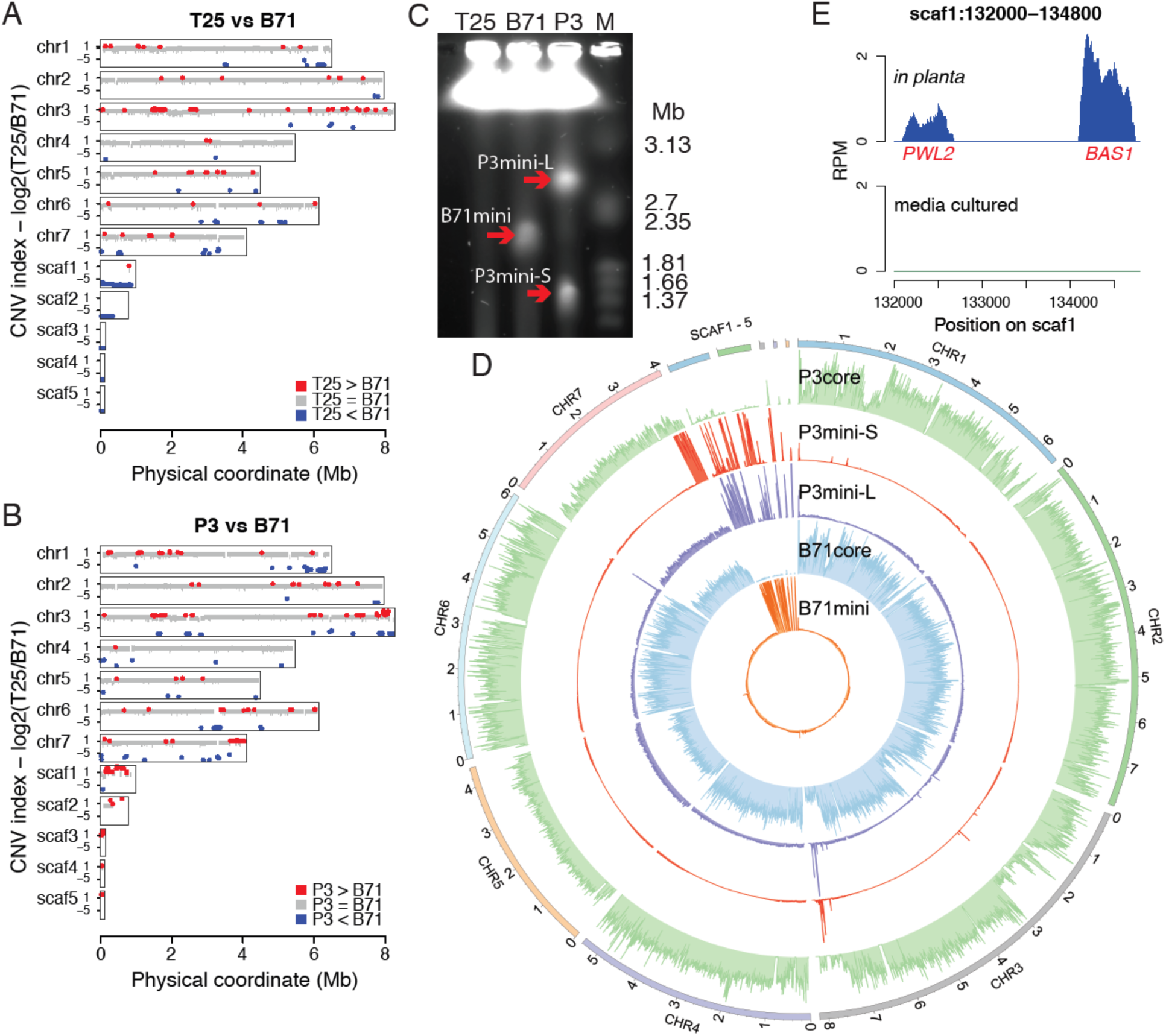
Genome comparison and mini-chromosomes. (**A, B**) Comparisons of T25 versus B71 (**A**) and P3 versus B71 (**B**). CNV index (y-axis) represents the log2 value of the ratio of read counts of a genomic segment between two isolates. Red, blue, gray lines represent CNplus, CNminus, and CNequal, respectively. (**C**) CHEF gel of three MoT isolates. Red arrows indicate mini-chromosomes. (**D**) Genome-wide distributions of read depths of 10kb bins, determined by uniquely-mapped reads, from sequencing of gel-excised core or mini-chromosomes. The 99.5% percentile of read depths per bin sets track height. (**E**) RNA-Seq reads distribution of *PWL2* and *BAS1*. RPM (reads per million of total aligned reads) represents normalized read counts.

### Dispensable mini-chromosomes of MoT strains

Variability in the five scaffolds led us to hypothesize that some or all scaffolds might correspond to a mini-chromosome in B71. Electrophoretic karyotypes of B71 using contour-clamped homogeneous electric field (CHEF) electrophoresis confirmed that B71, indeed, contained a mini-chromosome of ~2.0 Mb in size (**Fig. 4C**). Mini- and core chromosomal DNAs were separately excised from the gel for Illumina sequencing. The five scaffolds were highly over-represented among reads obtained from the mini-chromosome DNA and highly under-represented among the core chromosome reads (**Fig. 4D**). The mini-chromosome contains 192 protein-coding genes. Gene ontology (GO) enrichment analysis identified a significantly over-represented GO term, cysteine-type peptidase activity (GO:0008234, p-value=0.0001). Eight out of all 11 genes associated with cysteine-type peptidase activity are located on the mini-chromosome (**Data S6**). Known effector genes *PWL2* and *BAS1,* which are located on different core chromosomes in MG8, were located immediately adjacent to one another on the B71 mini-chromosome (**Fig. S4A**). This configuration was supported by 211 PacBio long reads and by Sanger sequencing of a PCR product obtained with a *PWL2* and *BAS1* primer pair (**Fig. S4B**). No *PWL2* or *BAS1* homologs, with at least 70% identity, were identified on core chromosomes. Both genes exhibited *in planta-* specific expression on the mini-chromosome (**Fig. 4E and Fig. S5**). Therefore, mini-chromosomes harbor effector genes that show similar *in planta*-specific expression patterns to effector genes residing on core chromosomes.

**Fig. 5.**
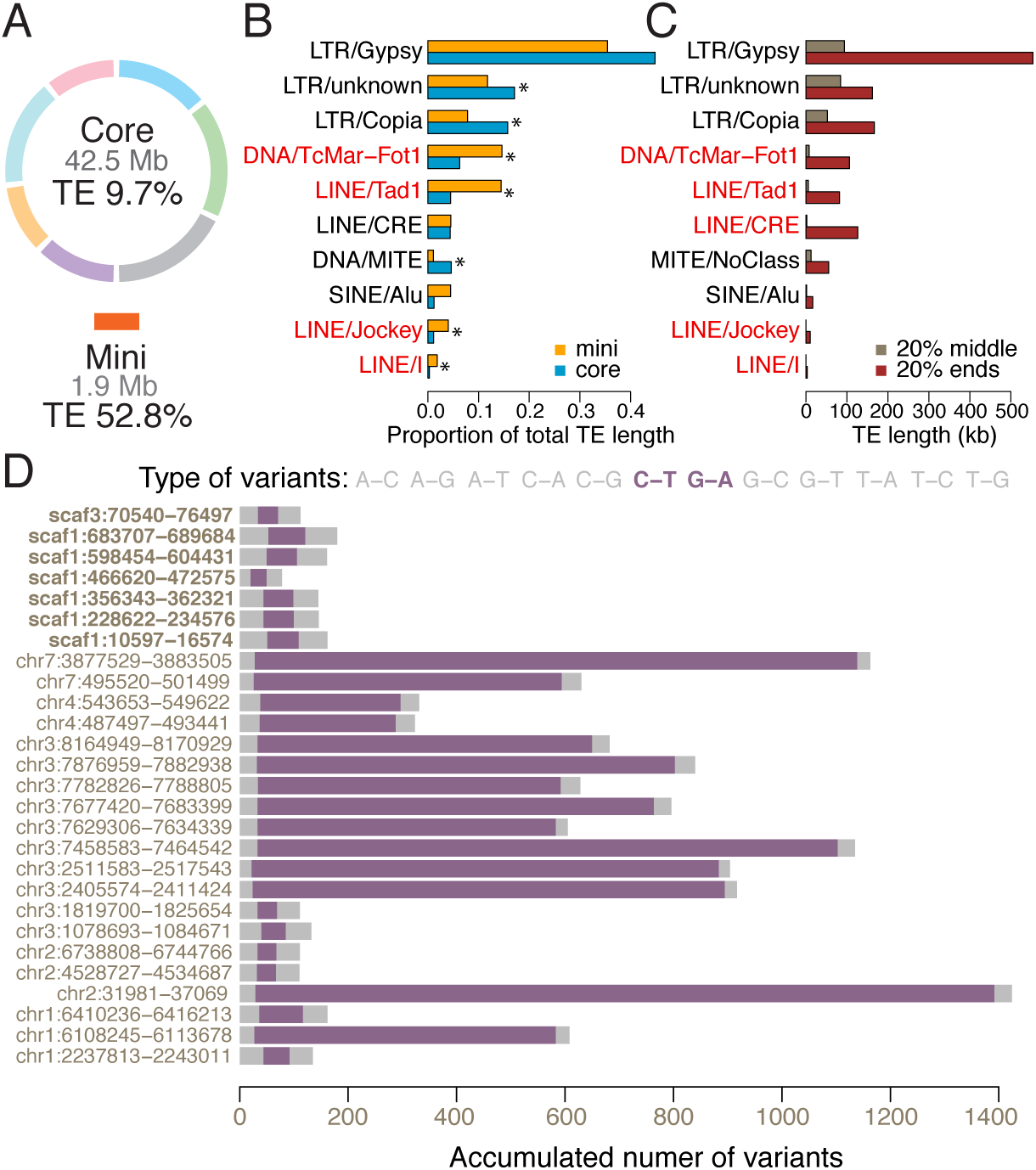
Comparison of repeats between core and mini-chromosomes. (**A**). Proportions of transposable elements (TE) in core and mini-chromosomes. (**B**) Barplots of proportions of each subclass out of total transposon sequences in core and mini-chromosomes. Subclasses with significantly proportional differences between core and mini-chromosomes are labeled with an asterisk (*), and subclasses over-represented in mini-chromosomes are highlighted in red. (**C**) Barplots indicate length of transposon subclasses in 20% ends and 20% middle regions of core chromosomes. Subclasses with at least 10-fold reduction in 20% middle versus 20% ends are highlighted in red. (**D**) Distribution of different variant types in MGR583 homologs relative to a reference MGR583 (e.g., A-C represents base A in reference MGR583 has changed to C in its homologs). RIP-type mutations are highlighted in purple. Name labels show the genomic coordinate of each homolog.

Further CHEF analyses showed no evidence of mini-chromosomes in T25 and supported two mini-chromosomes in P3, consistent with predictions from the CNV results. The P3 mini-chromosomes are ~1.5 Mb and ~3 Mb in length (**Fig. 4C**). Sequences of both P3 mini-chromosomes exhibited similarities to the B71 mini-chromosome but also marked differences (**Fig. 4D and Fig. S6**). The large P3 mini-chromosome contained both *PWL2* and *BAS1* genes (**Fig. S7**), plus it harbored ~33 kb (assembly location 6,007 to 6,039 kb) of duplicated DNA from a region near the end of chromosome 6, which included a homolog of the MoO effector *AVR-Pib* (Zhang *et al.* 2015). In contrast, the small P3 mini-chromosome lacked the *PWL2* and *BAS1* genes, but it contained a duplication of approximately 0.39 Mb of the chromosome 7 end (assembly location ~3.65 to 4.04 Mb) (**Fig. 4D**). Retention of this segment in the core chromosome explains the large CNplus segment at this region of P3 chromosome 7 (**Fig. 4B**). Notably, this segment contained five putative effector genes and a homolog of the known MoO effector gene *AVR-Pik* (Kanzaki *et al.* 2012). Another notable region from the end of chromosome 3 was present in both P3 mini-chromosomes, but not present in the B71 mini-chromosome (**Fig. 4D**). Sequencing P3 core chromosomes identified sequences homologous to the B71 mini-chromosome that were not present in B71 core chromosomes (**Fig. 4D**). Taken together, these three MoT mini-chromosomes contain different sets of known or predicted effector genes and other core-chromosome end sequences, which are either missing or duplicated on the core chromosomes of the same or other strains. The highly variable structure of MoT mini-chromosomes indicates frequent acquisition of sequences from core chromosomal ends.

### Repetitive sequences in B71 core- and mini-chromosomes

Repeat annotation showed approximately 12.9% of the B71 genome consisted of transposons and other repetitive elements, and transposons accounted for 9.7% and 52.8% of the core and mini-chromosomes, respectively (**Fig. 5A and Table S5**). Many of the transposons that were over-represented in the mini-chromosome occurred frequently on chromosome arms, particularly at chromosome ends (**Fig. S8**). Four transposon subclasses made up a greater proportion of the total transposon sequences on the mini-chromosome versus core chromosomes, including three LINEs (Tad1, Jockey and I) and the DNA transposon TcMar-Fot1 (**Fig. 5B**). These four are among the top five elements enriched in the core chromosomal 20% ends relative to the 20% middle core chromosome regions (**Fig. 5C**). Besides similarities in transposon composition between chromosome ends and the mini-chromosome, alignment of the B71 mini-chromosome sequence to core chromosomes identified duplications of >10 kb fragments with at least 95% identity. Duplications were located at ends of chromosomes 3, 4, and 7 (**Fig. S8**), and they were highly enriched for telomere-associated MoTeRs (LINE/CRE element). Therefore, a subset of MoT transposons is implicated in dynamic interactions between MoT mini-chromosomes and core chromosome ends.

Nucleotide composition analysis indicated that, overall, repetitive sequences along core chromosomes were highly negatively correlated with GC content (**Fig 3B, 3C**, **Fig. S9**). However, the highly negative correlation did not hold in the mini-chromosome, which is highly repetitive while maintaining relatively high GC content. Repetitive sequences in many fungi are subject to repeat-induced point (RIP) mutation resulting in C-to-T or G-to-A transitions and, thereby, leading to reduced GC content (Nakayashiki *et al.* 1999; Ikeda *et al.* 2002; Hane and Oliver 2008; Gladyshev 2017). Given higher GC content of repetitive sequences in the mini-chromosome versus core chromosomes, we explored the possibility of different levels of RIP in these genomic regions, as observed in *Fusarium* mini-chromosomes (Vanheule *et al.* 2016), by assessing their RIP-type mutation rates. Of six high-abundance transposons examined, all exhibited reduced levels of RIP-type mutations in the mini-chromosome relative to core chromosomes (**Fig. S10**). We examined transposons MGR583 (LINE/Tad1 element) and Pot2 (DNA/TcMar-Fot1 element) with multiple copies in both core and mini-chromosomes. RIP analysis indicated that no sequences of MGR583 (N=7) or Pot2 (N=22) from the mini-chromosome were subjected to extensive RIP-type mutations, while 14/20 MGR583 and 3/19 Pot2 from core chromosomes contained abundant RIP-type mutations (**Fig. 5D and Fig. S11**). Therefore, unlike transposons in core chromosomes, transposons in MoT mini-chromosomes do not appear to be inactivated by the RIP genome defense mechanism.

## Discussion

The B71 reference genome for the wheat blast fungus has shown a high degree of macrosynteny for the core chromosomes relative to the rice pathogen reference genome 70-15 (MG8), which supports the recent report maintaining *M. oryzae* as a single species (Gladieux *et al.* 2018). In contrast, mini-chromosomes present in B71 and another recent MoT field isolate P3 (P3-large and P3-small mini-chromosomes) are highly variable, with each one containing shared and different MoO effector homologs, putative effector genes, and other sequences from core chromosome ends. The B71 and P3-large mini-chromosomes contain the only copies of known MoO effectors *PWL2* and *BAS1* in these strains and neither gene was present in the early MoT strain T25, which lacks mini-chromosomes. *PWL2* and *BAS1* are located on different core chromosomes in 70-15, but they are found side-by-side on the B71 mini-chromosome. Both effectors show similar *in planta* specific expression on the MoT mini-chromosomes and on the MoO core chromosomes. Only the P3-large mini-chromosome contains a homolog of the MoO *AVR-Pib* gene, and only the P3-small mini-chromosome contains a homolog of *AVR-Pik*. Each mini-chromosome contains many other sequences that are either duplicated from core chromosome ends or missing from core chromosomes altogether. In one case, a P3 core chromosome sequence was homologous to the B71 mini-chromosome but not present in B71 core chromosomes. Taken together, our findings provide new insight on the two-speed genome of *M. oryzae* (Raffaele and KAMOUN 2012). That is, in contrast to housekeeping genes in slowly evolving core chromosome regions, *M. oryzae* effectors often occur in rapidly evolving, transposon-rich regions near chromosome ends. We expand understanding of the rapidly-evolving effector compartment in *M. oryzae* to include two apparently interchangeable regions, non-dispensable core chromosome ends coupled to dispensable mini-chromosomes.

The mechanism for sequence exchange between core- and mini-chromosomes is unknown. However, the enrichment in mini-chromosomes of multiple subclasses of LINE retro-transposons and a DNA transposon that are also enriched at core chromosome ends, points to a transposon-mediated recombination mechanism involving non-allelic homology. Such a mechanism has been shown to facilitate genome rearrangements in another phytopathogenic fungus (Faino *et al.* 2016). In contrast to seemingly RIPed core chromosome copies, the multiple copies of both MGR583 (LINE element) and Pot2 (DNA element) in the mini-chromosomes are nearly devoid of RIP-type mutations. This suggests that transposons on the mini-chromosomes remain active, facilitating multiplication and recombination. Telomere-associated MoTeR elements, found in MoL strains but not in MoO strains, are enriched in duplicated sequences on MoT mini-chromosomes. MoTeR elements have been reported to account for the extreme sequence variability of MoL telomeres compared to MoO telomeres (Starnes *et al.* 2012), suggesting these elements might enhance mini-chromosome dynamics in MoT and MoL strains through destabilization of telomere regions. Transposon-rich genomic regions have been linked to increased sequence and structural variation in fungal plant pathogens (Raffaele and Kamoun 2012; Yoshida *et al.* 2016; Moller and Stukenbrock 2017). Therefore, transposon-rich mini-chromosomes that also carry a number of genes, including many putative effectors, likely serve as genomic hotspots promoting genomic variation. Exceptional genomic variation produced in mini-chromosomes, and capable of flowing into core chromosomes, could accelerate the evolutionary potential of the pathogen.

Dynamic interchange between mini-chromosomes and core chromosome ends would explain *AVR* effector gene dynamics described in rice pathogens, which are notorious for their rapid ability to overcome deployed *R* genes. Corresponding *AVR* genes are most often deleted from MoO strains in response to *R* gene deployment (Valent and Khang 2010). Numerous *AVR* genes reside near telomeres in MoO field isolates and a couple have been localized to mini-chromosomes (Orbach *et al.* 1996; Luo *et al.* 2007; Chuma *et al.* 2011). The best understood case is *AVR-Pita1*, which confers avirulence to rice carrying the *Pita* gene (Khang *et al.* 2008). *AVR-Pita1* belongs to a telomeric gene family and shows a high rate of spontaneous mutations, including frequent deletions (Valent and KHANG 2010). The *AVR-Pita1* gene has also been highly mobile in the *M. oryzae* genome, having undergone multiple translocations to at least 8 distinct loci, usually near telomeres, across 6 different chromosomes including mini-chromosomes (Chuma *et al.* 2011). Mini-chromosomes would provide a population-wide repository for *AVR* genes that are deleted from individual strains and a means for rapid loss of *AVR* gene function from individual strains, because mini-chromosomes are frequently lost during vegetative growth and in the sexual cycle (Orbach *et al.* 1996). Individual strains lacking *AVR-Pita1* could regain it through acquiring *AVR-Pita1* containing mini-chromosomes from other individuals through the parasexual cycle and lateral gene transfer (Chuma *et al.* 2011), with the gene integrated into new locations on the core chromosomes, most likely at chromosome ends. The dynamic coupling we report between mini-chromosomes and core chromosome ends supports the multiple translocation hypothesis for *AVR* genes responding to periodic negative selection pressure of *R* gene deployment.

Collectively, we propose that the mini-chromosome plays a role for gene movements like a shuttle, in which mutation, duplication, loss, and rearrangements of DNA occur at a faster pace than normal genomic changes, hence, accelerating genomic evolution for adaptation. Our results together with the B71 reference genome will facilitate answering important questions for blast on wheat and other cereal crops. It is critical to monitor the evolution, including potential recombination with other *M. oryzae* pathotypes, of the complex MoT population in South America and the initially clonal MoT population in South Asia (Malaker *et al.* 2016; Cruz and Valent 2017). Localization of the *PWL2* host species-specificity gene on mini-chromosomes in wheat pathogens raises the question of a role for mini-chromosomes in host jumps. Effector gene dynamics, so far only associated with a small number of MoO *AVR* effector genes corresponding to periodically deployed *R* genes, raises the question of what roles known MoO *AVR* effector homologs and *BAS1* (lacking known *AVR* activity in MoO strains) play in wheat infection by MoT strains. Finally, it is critical to identify and deploy effective wheat blast resistance.

## Materials and Methods

Detailed description of materials and methods is included in SI Materials and Methods.

### Genetic materials

All *M*. *oryzae* strains examined were field strains from South America (**Table S1**). MoT isolates B71, T25, and P3 were isolated in Bolivia (2012), Brazil (1988), and Paraguay (2012), respectively. All work with living wheat blast fungus in the U.S. was performed with proper USDA-APHIS permits and monitoring in BSL-3 laboratories in the Biosecurity Research Institute at Kansas State University.

### DNA/RNA extraction

Single spore isolates of each pathogen strain were cultured on complete medium for mycelium propagation. Mycelium was harvested and frozen using liquid nitrogen. To avoid excessive mitochondrial DNA, mycelial nuclei were collected by gradient centrifugation as described (Zhang *et al.* 2012). The CTAB (cetyltrimethylammonium bromide) DNA extraction method was applied to isolate genomic DNA from the nuclear samples (Clarke 2009). RNA was extracted using RNeasy Plant Mini Kit.

### B71 genome sequencing and assembly

The 3-20 kb WGS libraries were constructed using B71 nuclear genomic DNAs. The library was sequenced with P6-C4 chemistry on ten SMRTcells of PacBio RS II. Nuclear genomic DNAs were also subjected 2×250 bp paired-end Illumina sequencing. To increase the assembly continuity, LIEP was devised and used to generate 20-30 kb long-distance paired sequences for scaffolding. PacBio long reads were assembled using the Canu pipeline (Koren *et al.* 2017). Self-correction using PacBio reads did not correct all PacBio sequencing errors. Illumina reads and the Illumina assembly sequences assembled using DISCOVAR de novo (Weisenfeld *et al.* 2014) were both utilized for further error correction. The resulting assembled contigs were scaffolded using LIEP long-distance paired sequences with the software SSPACE (Boetzer *et al.* 2011).

### Genome annotation

A Maker pipeline was used for the B71 genome annotation (Cantarel *et al.* 2008). Both evidence-driven prediction and *ab initio* gene prediction were employed (SAlamov and SOlovyev 2000). Transcriptional evidence was provided using assembled sequences from RNA sequencing data of the B71 strain that was cultured in media and field wheat leaf samples infected by Bangladesh wheat blast strains, which were genetically close to B71. CEGMA was used to assess the completeness of the genome assembly or annotation (Parra *et al.* 2009).

### Identification of putative effectors

Publically available RNA-Seq data of MoT infected wheat were used as *in planta* expression data to compare with RNA-Seq data from a cultured sample. Genes with read abundance higher than 1 FPKM (fragment per kilobase of coding sequence per million reads) from the *in planta* data set but no reads from the cultured sample were considered to be *in planta* specific expression. *In planta* specific genes containing classical signal peptide domains (Nielsen 2017) were considered putative effectors.

### Analysis of copy number variation between strains

Read depth approach was employed to identify CNV between each of some *M. oryzae* strains and B71 for each of sequence bins (e.g, 300 bp). Segmentation with the R package of DNACopy was performed to identify genomic CNV segments merged from multiple bins (Olshen *et al.* 2004).

### CHEF karyotypes of MoT strains and mini-chromosome sequencing

MoT protoplasts were prepared and mixed with 1.5% low melting-temperature agarose (Orbach *et al.* 1996). Suspensions were loaded into disposable plug molds. Protoplasts in plugs were lysed with proteinase K and washed. A Biorad CHEF electrophoresis system was used for separating chromosomes embedded in the plugs. After the CHEF gel electrophoresis, DNAs from individual mini-chromosomes, and from core chromosomes as one unit, were excised and purified from the agarose gels. Purified DNAs were subjected to Illumina 2×151 bp paired-end sequencing.

### Analyses of repetitive sequences

Repetitive sequences were identified using MGEScan (Lee *et al.* 2016), LTR_Finder (Xu and Wang 2007), LTRharvest (Ellinghaus *et al.* 2008; Gremme *et al.* 2013), and RepeatModeler (github.com/rmhubley/RepeatModeler). Merging discovered repetitive sequences and previously characterized *M. oryzae* repeats (Bao *et al.* 2015) produced a non-redundant database, which served as a repeat library to identify repeats in the B71 genome using RepeatMasker (www.repeatmasker.org). Some transposable elements were subjected to analysis of RIP-type polymorphisms, nucleotide changes of C to T or G to A.

## Acknowledgements

Data are available from Sequence Read Archive: PRJNA355407. This project has been funded by Agriculture and Food Research Initiative Competitive Grant #2013-68004-20378 from the USDA National Institute of Food and Agriculture. We thank Drs. Nicholas J Talbot and Yukio Tosa for critical review prior to manuscript submission and thank Dr. Alina Akhunova at the Integrated Genomics Facility at Kansas State University for technical support. Photos in Figs. 1B and 1C are from Guillermo Isidoro Barea Vargas, Coperagro SRL, Bolivia. The photo in Fig. 1D is from Javier Toledo, Agripac Boliviana, Bolivia. This is the contribution number 18-336-J from the Kansas Agricultural Experiment Station.

## Supplementary Information (SI)

Effector Gene Reshuffling Involves Dispensable Mini-chromosomes in the Wheat Blast Fungus

Zhao Peng, Ely Oliveira Garcia, Guifang Lin, Ying Hu, Melinda Dalby, Pierre Migeon, Haibao Tang, Mark Farman, David Cook, Frank F. White, Barbara Valent, Sanzhen Liu

## Materials and Methods

### Pathogen strains and Biosafety-Level 3 (BSL-3) research

Fungal strains used are described in **Table S1**. MoT strain B71 was isolated from wheat field in Bolivia in 2012. MoT isolates T25 and P3 were collected from Brazil and Paraguay in 1988 and 2012, respectively. All work with living wheat blast fungus was performed in BSL-3 laboratories in the Biosecurity Research Institute at Kansas State University and in Foreign Disease-Weed Science Research Unit at Fort Detrick in Maryland. The strains are permanently stored in a collection maintained at Fort Detrick.

### DNA extraction

Single spore isolation of each MoT strain was performed to ensure genetic purity before DNA isolation. The strains were cultured on oatmeal or rice polish agar plates. The fungal culture was then transferred to complete medium for mycelium propagation at room temperature in a shaker. Mycelium was harvested by passing the fungal culture through a funnel covered with sterile filters. The mycelium was then frozen using liquid nitrogen and stored at −80°C. To avoid excessive mitochondrial DNA, mycelial nuclei collected by gradient centrifugation following protocol described by (Zhang *et al.* 2012). The CTAB (cetyltrimethylammonium bromide) DNA extraction method was applied for nuclear samples to isolate genomic DNA (Clarke 2009).

### PacBio sequencing

The 3-20kb whole genome shotgun libraries were constructed using nuclear genomic DNAs of the B71 strain. The library was sequenced with P6-C4 chemistry on ten SMRTcells of PacBio RS II at the Yale Center for Genomic Analysis (YCGA).

### Illumina sequencing

Nuclear genomic DNAs of the B71 strain were subjected to library preparation using a TruSeq PCR-free library protocol with the insertion size of 375 bp. 2×250 bp paired-end reads were generated on an Illumina HiSeq2500 at Beijing Genomics Institute.

### Construction of indexed LIEP vectors

Two synthetic sequences with random barcodes and Illumina compatible sequence were annealed, end-filled and cut by *Mfe*I and *Hind*III restriction enzymes (NEB, USA) (**Fig. S12**). DNA fragments with random barcodes were referred to as linkers. Linkers with random barcodes were then inserted into the commercialized vector pEZ-BAC (Lucigen, USA) digested by *EcoR*I and *Hind*III (NEB, USA). Three millions of clones with random barcoded vectors were pooled.

### Linker sequencing of LIEP barcode-indexed vectors

Plasmid DNAs of barcode-indexed vector were extracted and Illumina adaptor primers were used to amplify barcodes for sequencing. Two steps of PCR amplification using Phusion High Fidelity DNA polymerase (NEB, USA) were applied. In the 1^st^ step PCR, the Illumina compatible primer RPtag and vector unique primer L95 (**Table S6**) was used, following the protocol: 98°C 30 seconds, 5 cycles of (98°C 10 seconds, 57°C 15 seconds, 72°C 15 seconds), and 72°C 2 minutes. After purification of the 1^st^ step PCR product, it was directly used in the 2^nd^ step PCR by using modified Illumina primers htF501s2 and htF701s (**Table S6**), following the same PCR protocol in first step except changing the T_m_ to 61°C. Libraries were sequenced on a MiSeq with 2×78 cycles at the Integrated Genomic Facility at Kansas State University and on a HiSeq2000 with 2×100 cycles at the Genome Sequencing Facility at the Kansas University Medical Center. Two barcodes of a barcode pair of each vector were extracted from each sequencing read using an in-house Perl script. Barcode data were kept only when identical barcode sequences were extracted from two paired reads. Unique barcode pairs were obtained after removing redundancy.

### LIEP library preparation and sequencing

The genomic DNA of B71 strain was partially digested by *Sau3A*I restriction enzyme (NEB, USA), around 20-30 kb DNA fragments were selected to ligate with *BamH*I digested barcoded vector pool, which had been dephosphorylated with rSAP (NEB, USA) to avoid self-ligation. Eight pools of BAC clones were obtained and individual pool contained 8,000-25,000 unique clones. In total, approximately 100,000 of BAC clones were obtained. The DNA of each BAC pool was extracted using regular plasmid preparation protocol. The regular Illumina library preparation protocol was modified for sequencing the BAC ends. The DNA of each BAC pool was sheared into 500-800bp fragments in Coravis ultrasonicator (Integrated Genomic Facility, Kansas State University). End repair and dA-tailing with ddATP (Sigma, USA) of the sheared DNA was performed following the NEB Next Ultra End repair/ dA-tailing Module. T-adaptor was prepared by annealing the hta1 and hta2 oligos (**Table S6**) under the following protocol: heating at 95°C for 5 minutes, cooling to 25°C at the rate 1°C every 15 seconds, and then holding at 25°C for 30 minutes in thermal cycler. A-tailing product and the T-adapter was ligated by T4 DNA ligase (NEB, USA). Two steps of PCR amplification using Q5 DNA polymerase (NEB, USA) were then applied. In 1^st^ step PCR, primers 501Tr1 and 701Tr1 (**Table S6**) were used, following the protocol: 98°C 1 minute, 12 cycles of (98°C 10 seconds, 66°C 15 seconds, 72°C 30 seconds), and 72°C 2 minutes. In the 2^nd^ PCR, different indexed primers from Illumina Truseq HT primer sets (**Table S6**) were applied for amplifying the 1^st^ PCR products, following the same PCR conditions but only 5 cycles. The resulting DNA libraries were pooled in ratios according to the size of BAC libraries, and sent for MiSeq sequencing at the Integrated Genomic Facility at Kansas State University.

### LIEP data analysis

For each BAC clone, the DNA insert was flanked by two indexed barcodes that were built in a vector. The barcodes were known based on previously vector linker sequencing. That means, based on barcode information, we can figure out a pair of two insert end sequences even through two ends were sequenced separately. The LIEP data analysis is to process sequencing data to extract barcode sequence and insert sequences, and finally determine two BAC end sequences for each clone. In detail, each LIEP raw sequence (PE reads) started with a barcode followed by the sequence of a genomic DNA insert. LIEP sequencing raw sequences were first subjected to adaptor trimming (trimmomatic-0.36), without quality trimming, using TruSeq adaptor sequences, followed by extracting indexed barcode sequences. After this step, each sequence had an associated barcode, which was then used for barcode trimming to avoid barcode contamination in sequences. Quality trimming was performed at the same time. The barcode was also used to identify the paired barcode from the same vector, which was pre-determined. For example, a pair of indexed barcodes, BC-I and BC-II, was identified. BC-1 associated reads represent sequences from one end of the insert of the BAC clone, while BC-2 associated reads represent sequences from the other end of the same BAC clone. Multiple reads of each end were obtained if a BAC clone was sequenced multiple times. We then assembled BC-1 and BC-2 associated reads separately with Celera Assembler 8.0 to obtain sequences of two BAC ends (Myers *et al.* 2000).

### PacBio genome assembly

Only PacBio reads longer than 5 kb were used for the assembly. Canu v1.3-r7616 was selected to correct reads, to trim suspicious sequences (e.g., adaptors), and to assemble corrected and cleaned reads into unitigs (Koren *et al.* 2017).

### Illumina genome assembly

Trimmomatic (version 0.36) was used to trim adaptor sequences of TruSeq PCR free 2×250bp sequencing reads (Bolger *et al.* 2014). Trimmed reads after removing adaptors were subjected to error correction with the error correction module of ALLPATHS-LG (GNERRE et al. 2011). Corrected reads were *de novo* assembled using DISCOVAR de novo (Weisenfeld *et al.* 2014). The final Illumina assembly contains contigs with at least 500 bp.

### Genome assembly polishing and scaffolding

The tools pbalign and Quiver were used to align PacBio reads to the PacBio assembly and polishing. Self-correction in Canu and Quiver using PacBio reads corrected most sequencing errors. However, a number of sequencing errors existed in the draft PacBio assembly, particularly on homopolymeric regions. Two rounds of additional polishing were employed to improve the assembly quality. First, trimmed clean PCR free Illumina genomic DNA sequencing reads were directly aligned to the draft PacBio assembly with the *mem* module of BWA (0.7.10-r789) (Li and Durbin 2010). Alignments were filtered using the following criteria: at least 100 bp matches with 94% identity, at least 98% coverage, and at least 40 of mapping score. Alignments passing the filtering criteria were used to call small INDELs (insertions and deletions) with GATK (version 3.3), followed by a set of filtering criteria (MCKenna *et al.* 2010). Each INDEL required at least 10 but no more than 5,000 reads supported and the non-reference variant (an alternative sequence type relative to the draft assembly) of each INDEL site accounted for at least 90% reads of the total supporting reads covering the variant site. The corrected draft assembly with this set of INDELs was obtained by using the consensus module of bcftools (Li 2011).

To reduce errors during the correction using short Illumina reads due to false alignments, long sequences of the DISCOVAR de novo assembly were utilized for the second round of correction. Assembled sequences were aligned to the first-round corrected assembly with the mem module of BWA (0.7.10-r789) (Li and Durbin 2010). Alignments were filtered using the following criteria: at least 400 bp matches with 94% identity, at least 98% coverage, and at least 40 of mapping score. Alignments passing the filtering criteria were used to call single nucleotide variants and small INDELs (insertions and deletions) with a pipeline using samtools (mpileup module) and bcftools (the module of “call”). Again the consensus module of bcftools was implemented the second round correction, resulting in a new draft assembly. Contigs of the draft assembly were scaffolded using LIEP BAC end sequences. The software SSPACE 3.0 was used for scaffolding with the parameter of “-x 0 -k 3 -g 1 bwasw end1.fq end2.fq 30000 0.8 FR” in that end1.fq and end2.fq represent sequence FASTQ files of first ends and second ends of BACs, respectively (Boetzer *et al.* 2011). The scaffolding required at least three pairs of LIEP sequences to establish a connection between two contigs.

### *De novo* RNA-Seq assembly

The mycelium grown in the complete medium, described in the DNA extraction, was also applied for RNA extraction using RNeasy Plant Mini Kit (Qiagen, USA). Total RNA was used to prepare an mRNA sequencing library that was run on a MiSeq with 2×150 cycles at the Integrated Genomic Facility at Kansas State University. The software Trimmomatic (version 0.36) was used to trim adaptor sequences of RNA sequencing reads. Only paired reads both of which are at least 50 bp after trimming were retained. Trimmed clean reads were *de novo* assembled using Trinity with the default parameter (Grabherr *et al.* 2011).

### Identification of putative effectors

RNA-Seq data of MoT infected wheat were downloaded from Sequence Reads Achive, SRA number from ERR1360178 to ERR1360193. All data were merged to a single data set that was used as *in planta* RNA-Seq data. Both *in planta* RNA-Seq data and RNA-Seq data from a cultured sample were aligned to the B71Ref1 with STAR (2.5.2a). With the annotation file, Cufflinks v2.1.1 was used to determine read abundance per gene. Genes with read abundance higher than 1 FPKM (fragment per kilobase of coding sequence per million reads) from the *in planta* data set but no reads from the cultured sample were considered to be only expressed *in planta* or *in planta* specific expression, and vice versa. Genes that were expressed specifically *in planta* and contained classical signal peptide domains were considered putative effectors.

### Genome annotation

A Maker pipeline was used for the B71 genome annotation (Cantarel *et al.* 2008). Both evidence-driven prediction and *ab initio* gene prediction were employed (Salamov and Solovyev 2000). The B71 genome was repeat masked using a species-specific repeat database, which was generated by using RepeatModeler (1.0.9) (repeatmasker.org/RepeatModeler). EST evidence was provided by using Trinity assembled sequences from RNA sequencing data of the B71 strain that was cultured in media and field wheat leaves inoculated by Bangladesh wheat blast strains, which were genetically close to B71. Protein sequences of *M. oryzae* 70-15 (MG8) were provided as protein homologs. Augustus, GeneMark, and Snap were used for *ab initio* gene prediction. Snap was trained by using predicted gene models from a prior Maker run without using Snap.

### CEGMA analysis

CEGMA (v2.5) was used to compare predicted genes with 248 Core Eukaryotic Genes (CEG) to assess the completeness of the genome or annotations (Parra *et al.* 2009).

### Signal peptide analysis

Signalp-4.1 was used predicted the presence of signal peptide cleavage sites and transmembrane segments in protein sequences of each gene (Nielsen 2017).

### Gene ontology (GO) analysis

GO enrichment analysis was performed to find GO terms over-represented in the genes on the mini-chromosome. The resampling method with 10,000 samplings was employed (Young *et al.* 2010).

### Genome-wide variant discovery, genotyping, and construction of phylogeny

Assemblies of Mo isolates were downloaded from GenBank or from previous studies were used for the discovery of polymorphisms between each strain and the B71 reference genome. First, assembly contigs were split into multiple sequences if a contig is longer than 10 kb. Second, all resulting sequences from a strain were aligned to the B71 reference genome with BWA (0.7.10-r789). Each alignment required at least a 100 bp, at least 96% identity, at most 4% tail that can not be aligned to the reference at the ends of sequences, and at least 40 for the mapping score. GATK (version 3.3) used alignments to discover SNPs and genotyped each strain. Only bi-allelic SNP sites with at least 10% minor allele frequency, 20-500 supported sequences were retained at genotyping data. Genotyping data at SNP sites were used to construct a phylogenetic tree using an R package APE (Paradis *et al.* 2004).

### Analysis of genomic structural variation

A read depth approach was employed to examine genomic structural variation, specifically copy number variation (CNV), among some MoT strains. First, genomic bins each of which contains a certain length of non-repetitive sequences were defined. In detail, single-copy 25-mers were extracted from the B71 reference genome with Jellyfish (Marcais and Kingsford 2011). These k-mers were mapped back to the reference genome through BWA mapping. The genome was then scanned to find non-overlapping bins with 200 unique single-copy k-mers. Bin sizes were ranked from smallest to largest and the top 1% largest bins were removed for further analysis.

Second, Illumina reads of each strain were aligned to the B71 reference genome and read depths were determined and normalized by using total aligned reads for every defined genomic bin. For each bin, the log2 read depth ratio was calculated between the non-B71 strain and the B71 strain (log2(non-B71/B71)), which was then used for the segmentation to identify a continuous genomic segments exhibiting similar values of log2(non-B71/B71) of bins with the R package of DNACopy (Olshen *et al.* 2004). The module smooth.CNA with the parameters of (smooth.region = 10, outlier.SD.scale = 4, smooth.SD.scale = 2, trim = 0.01) and the module of segment with the parameter of (alpha=0.005, nperm=10000, p.method=“perm”, eta=0.005, min.width=3, undo.splits = “sdundo”, undo.SD = 3) were employed for the segmentation.

For a segment, the mean of log2(non-B71/B71) is close to zero if sequences of two strains are identical and no CNVs. The derivation of the mean of log2(non-B71/B71) from zero is due to sequence polymorphisms and copy number variation (CNV), including presence and absence variation (PAV). Thus, means of log2(non-B71/B71) of segments were used for the classification of segments to different categories. The classification also utilized information of read alignment coverage of segments determined by using Bedtools (the coverage module). For each segment, an alignment coverage ratio (relative coverage) was calculated by dividing the coverage of reads from a non-B71 strain to that from the B71 stain. A segment was categorized to CNVplus if the relative coverage is greater than 0.85 and the mean of log2(non-B71/B71) is greater than 0.6; segment was categorized to CNVplus segments if the relative coverage is greater than 0.85 and the mean of log2(non-B71/B71) is greater than 0.6; a segment was categorized to conserved segments, CNequal, if the relative coverage is greater than 0.85 and the mean of log2(non-B71/B71) is between −1.5 and 0.5; a segment was categorized to polymorphic segments if the relative coverage is between 0.2 and 0.8, and the mean of log2(non-B71/B71) is between −5 and −2; a segment was categorized to CNVminus segments if the relative coverage is less than 0.2, and the mean of log2(non-B71/B71) is less than −5. The mean of log2(non-B71/B71) of a segment was referred to as a CNV index of the segment.

### Electrophoretic karyotypes of MoT strains

Protoplasts of MoT strains were prepared following protocol described by Orbach et al. with some modification (Orbach *et al.* 1996). After resuspending fungal mycelia in 1M sorbitol (Sigma, USA), lysing enzyme (Sigma L-1412), instead of Novozym 234, was used to generate protoplasts. Purified protoplast was resuspended in SE buffer (1M sorbitol, 50mM EDTA, pH=8.0) at a density of 3×10^9^/ml. The protoplast solution was then mixed with 1.5% low melting-temperature agarose at a ratio of 1:1 volume. The mixture was loaded into disposable plug mold (Bio-rad, USA) to be solidified for 30 minutes on ice. The plugs were then lysed with proteinase K (DNAse free, Promega, USA) in the NDS buffer in a shaking water bath at 50°C overnight (Orbach *et al.* 1988). The plugs were washed with 50mM EDTA (pH=8.0) buffer at least three times at room temperature and then stored at 4°C. CHEF electrophoresis system (Bio-rad, USA) was applied for separating chromosomes DNA embedded in the plugs. The gels were made with 0.7% Certified Megabase agarose (Bio-rad, USA) and run in 0.5×TBE in the cold room under electrophoresis conditions as described (Xue *et al.* 2012).

### Sequencing of mini-chromosomes of MoT strains

After the CHEF gel electrophoresis, agarose containing B71 and P3 mini-chromosome DNAs was excised and transferred to D-tubes (Midi size) for electroelution following the provided manual (Millipore, Sigma, USA). The DNA was then concentrated by vacuum evaporation and dialyzed with distilled water on the Nitrocellulose membrane (Millipore, Sigma, USA) to remove salt. Meanwhile, B71 and P3 core chromosomes were excised as well and purified by Qiagen gel purification kit. Illumina TruSeq libraries were constructed with the purified DNA. Libraries were sequenced with 2×151 bp on a MiSeq in Integrated Genomic Facility at Kansas State University.

### Genomic PCR amplification of *PWL2* and *BAS1*

Using the genomic DNA of strains B71, P3 and T25 as templates, genes were amplified by Q5 high-fidelity DNA polymerase (NEB, USA) with primer pairs of MgActinF + MgActinR, BAS1-F + BAS1-R, and PWL2-F + PWL2-R (**Table S6**). Thermocycling conditions included initial denaturation at 98°C for 2 min, 25 cycles of 98°C for 8 s, 63°C for 30 s and 72°C for 30 s, and final extension at 72°C for 5 min.

### qRT-PCR

Infected wheat leaf tissues were obtained from 9-day old wheat seedlings (c.v. Bob White), inoculated with B71 (5×10^4^ spores/mL) and harvested 36 hours post inoculation. B71 mycelia were obtained from liquid cultures (3-3-3 media) grown in flasks on a rotating *shaker (100 rpm; 25 °*C) for 5 days. Total RNA from mycelium and infected wheat leaf tissues were extracted using miVana RNA isolation kit (Invitrogen) and pretreated with DNAse I amplification grade (1 U/µl) (Invitrogen Inc, Carlsbad, CA) according to manufacturer’s instructions. RNA quantity and quality were assessed, and cDNA was synthesized using 1 µg of total RNA extracted from each sample. RNA was reverse transcribed with 1 mM of Oligo (dT)_20_, 0.5 mM dNTP mix and 1 µL Superscript III reverse transcriptase (200 U/µl) (Invitrogen, Carlsbad, CA) in a final volume of 20 µL following manufacturer’s instructions. qRT-PCR was performed in a total volume of 10 µL, with 100 ng of total cDNA, using the IQ^TM^ SYBR^®^Green Super Mix reagent (BioRad, Hercules, CA) according to manufacturer instructions. The *M. oryzae* actin gene (MGG_03982) was amplified using MgActinF and MgActinR primers. Primers MgPwl2F and Pwl2_qRT2-R4 were used to amplify the *PWL2* gene. Primers MgBas1F and MgBas1R were used to amplify the *BAS1* gene. Melting curve and agarose gel analyses confirmed amplification of a single product. Cycle threshold values (*C_t_*) of three independent replicates were used to quantify gene expression. Relative expression (RR) levels of each gene of interest (goi) was calculated and normalized using the *actin* gene as reference (Liu *et al.* 2009; Mosquera *et al.* 2009) (*RR* = 100×2 ^(*c_t_actin*–*c_t_goi*)^). For **C*_t_* values of NA (Not Available), the value of 40 cycles was used.

### Identification of transposable elements

Transposable element DNA was identified using a combination of programs. Repeats were identified *de novo* using MGEScan (Lee *et al.* 2016), LTR_Finder (Xu and Wang 2007) and the LTRharvest program from GenomeTools (Ellinghaus *et al.* 2008; Gremme *et al.* 2013). Consensus Long-Terminal Repeat retrotransposons (LTR-RT) were identified from the output of these three programs using LTR_retriever (Ou and Jiang 2018). Putative Non-LTR retrotransposons identified from MGEScan were validated using BLASTx against a database containing reverse-transcriptase (RT) and transposase (DDE) domains. Elements containing significant similarity to either the RT or DDE1 domain, but not both, were kept and classified as either Long Interspersed Nuclear Elements (LINE) or a DNA transposon. Miniature Inverted-repeats were identified using MITE-hunter (Han and Wessler 2010). Putative MITEs containing multiple insertions in the genome were manually checked for the presence of terminal inverted repeat (TIR) domains and nucleotide divergence flanking either side of the elements. The LTR, LINE, DNA transposon and MITE elements were combined into a single database (intermediate DB) and clustered using the CD-HIT software. The resulting non-redundant database was used to mask repeats in the B71 assembly with RepeatMasker (Smit AFA, 2013-2015). An additional round of *de novo* repeat identification was carried out on the masked genome using RepeatModeler (Smit AFA, 2008-2015). Previously characterized repeats corresponding to *M. oryzae* from the RepBase database were also identified (Bao *et al.* 2015). The intermediate DB was combined with the RepeatModeler and RepBase repeats, and again clustered to produce a non-redundant database, which was served as a custom repeat library to identify repeats in the B71 genome using RepeatMasker.

### Analysis of RIP-type polymorphisms

Based on the RepeatMasker result, genomic sequences of the most abundant transposon elements were extracted and aligned to corresponding transposon sequences from the RepeatMasker database as the reference sequences. The GC percentages of the reference sequences were from 41.3% to 59.3%, which were used to examine relative RIP levels of queries. Only alignments with >60% coverage of a query that is the sequence from the genome were retained. Polymorphisms of each sequence were extracted from the alignments and types of mismatches were determined. The nucleotide changes of C-T or G-A from the reference sequence to the query were considered RIP-type variants. For each transposon sequence, the null hypothesis of that the mean of proportions of RIP-type variants out of the total mismatches transposons located at core chromosomes was not different with that of transposons located at the mini-chromosome was statistically tested using t-test.

For two long transposon elements, MGR583 and POT2, intact homologs were extracted from core chromosomes and the mini-chromosome. The levels of RIP-type mutations of individual sequences were also determined.

---

**Fig. S1.**
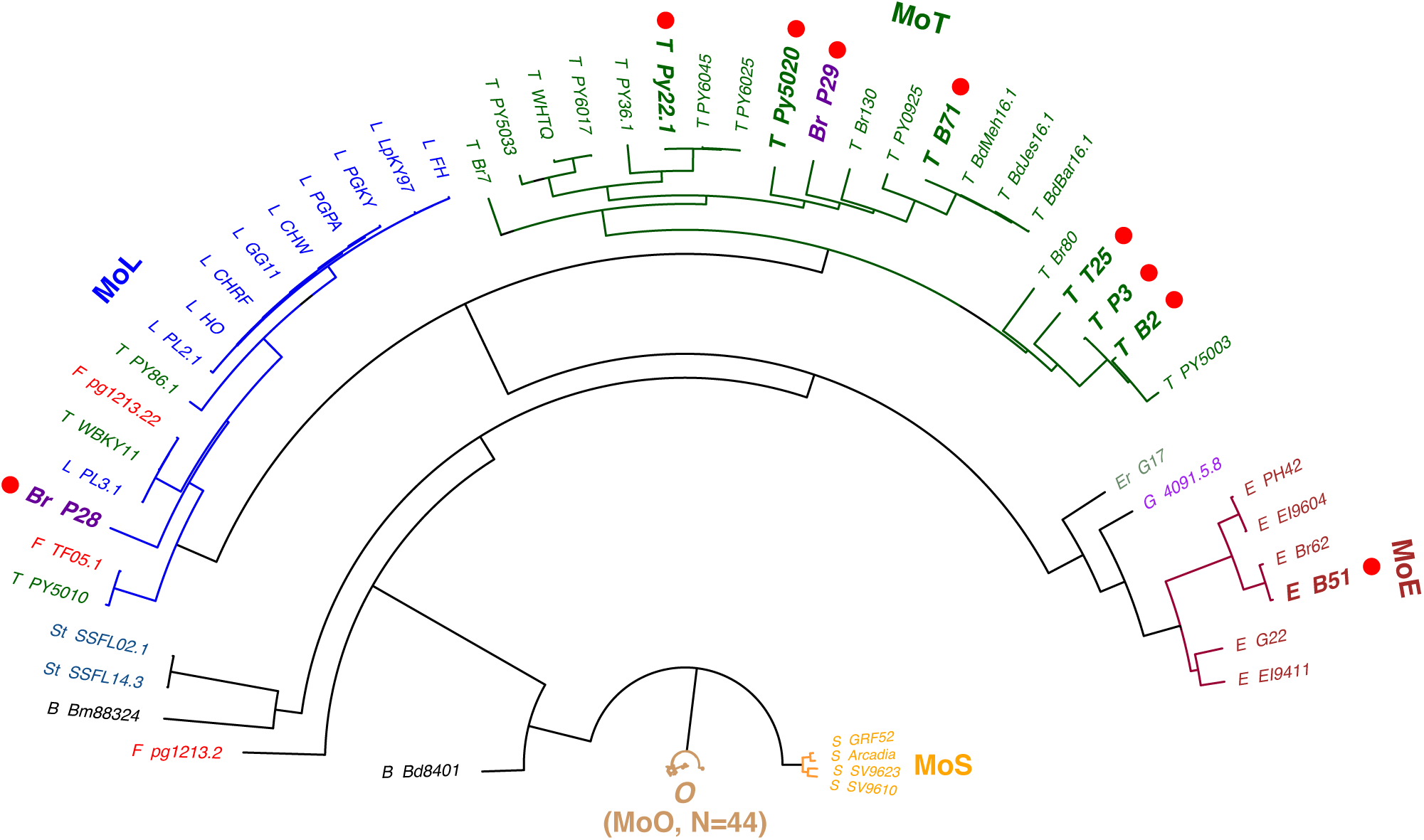
Phylogenetic tree of *M. oryzae* strains showing the major crop-specific pathotypes. These are: *Oryza* pathotype (MoO, 44 Strains); *Setaria* pathotype (MoS, 4 strains); *Eleusine* pathotype (MoE, 6 strains); *Triticum* pathotype (MoT, 21 strains); and *Lolium* pathotype (MoL, 16 strains) (Tosa *et al.* 2004; Farman *et al.* 2017). Strain branches in each of five pathotypes were labeled with the same color as the pathotype identifier. Assembly data of each strain were utilized to identify polymorphisms and construct the phylogeny with the neighbor-joining tree estimation. Strains selected in this study are highlighted with red dots. Host species on which each strain was isolated from the field are indicated (e.g., T) by: B, *Brachiaria*; Br, *Bromus*; E, *Eleusine*; Er, *Eragrostis*; F, *Festuca*; L, *Lolium*; O, *Oryza*; S, *Setaria*; St, *Stenotaphrum*; T, *Triticum*. The strain G 4091-5-8, which infects both *Eragrostis* spp. and *Eleusine* spp., was obtained in a laboratory cross between E G22 and Er G17. Strains Py22.1 and Py5020 are described in Pieck *et al*, 2017 (Pieck *et al.* 2017); and all other non-MoO strains are described in Gladieux *et al*, 2018 (Gladieux *et al.* 2018).

**Fig. S2.**
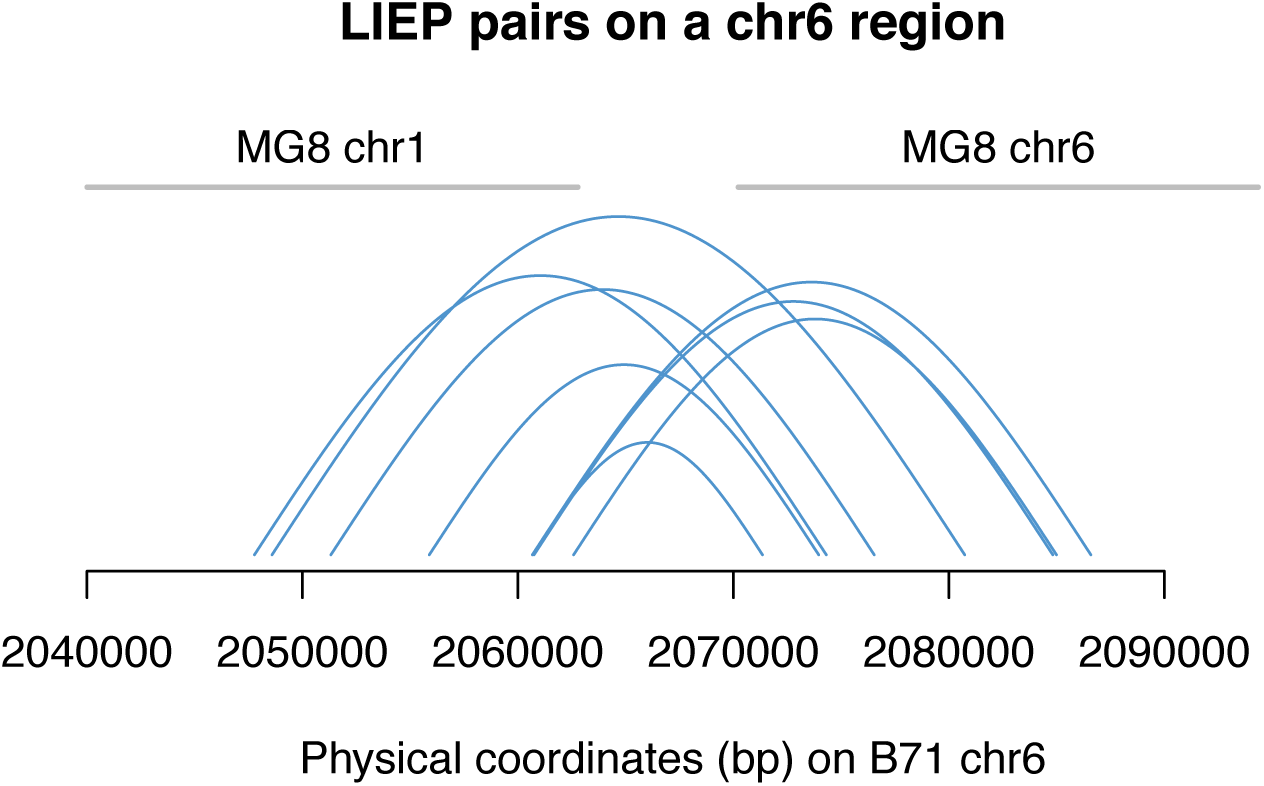
LIEP paired sequences on the rearrangement region on chromosome 6. The region (chromosome 6, 1-2,062,779 bp) of B71Ref1 is collinear with a partial sequence of MG8 chromosome 1, and the region beyond 2,070,228 bp of B71Ref1 chromosome 6 is collinear with MG8 chromosome 6. Blue curves showed pairs of LIEP sequences spanning the junction region, from 2,062,779 bp to 2,070,228 bp. In addition, the junction region and some flanking sequences are fully covered by 50 single PacBio long reads.

**Fig. S3.**
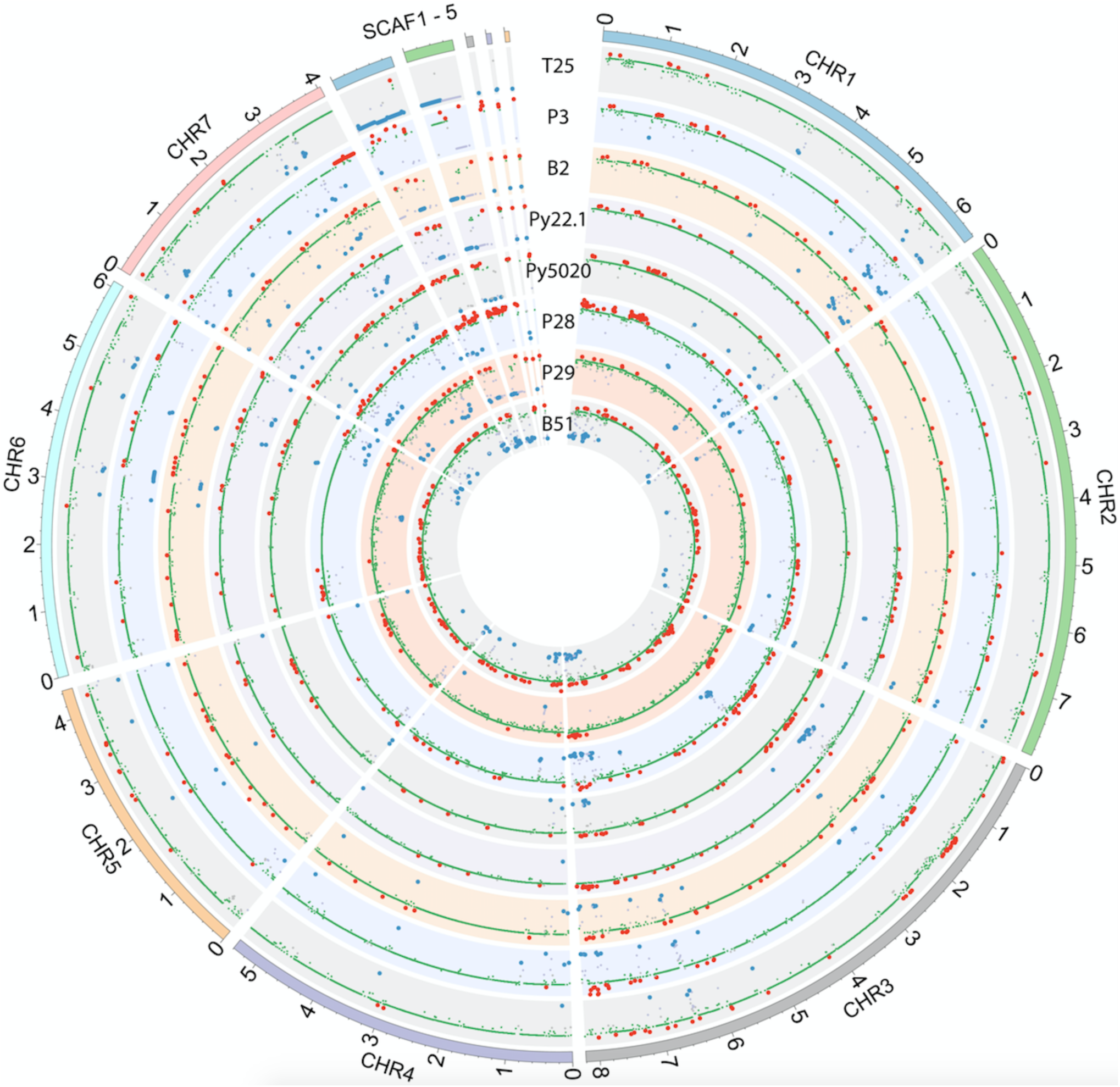
Genome comparisons between each of 8 additional *M. oryzae* strains and B71. The strains being compared are MoT strains T25, P3, B2, Py22.1, Py5020 and P29; the MoL strain P28; and the MoE strain B51 **(Fig. S2 and Table S1).** Each track represents a copy number comparison of the non-B71 isolate versus B71. The value of CNV index, which represents the log2 value of the ratio of sequencing read counts in genomic segments between two isolates of the comparison, determines vertical position on the track. Red, blue, green lines represent CNplus, CNminus, and CNequal regions relative to the B71.

**Fig. S4.**
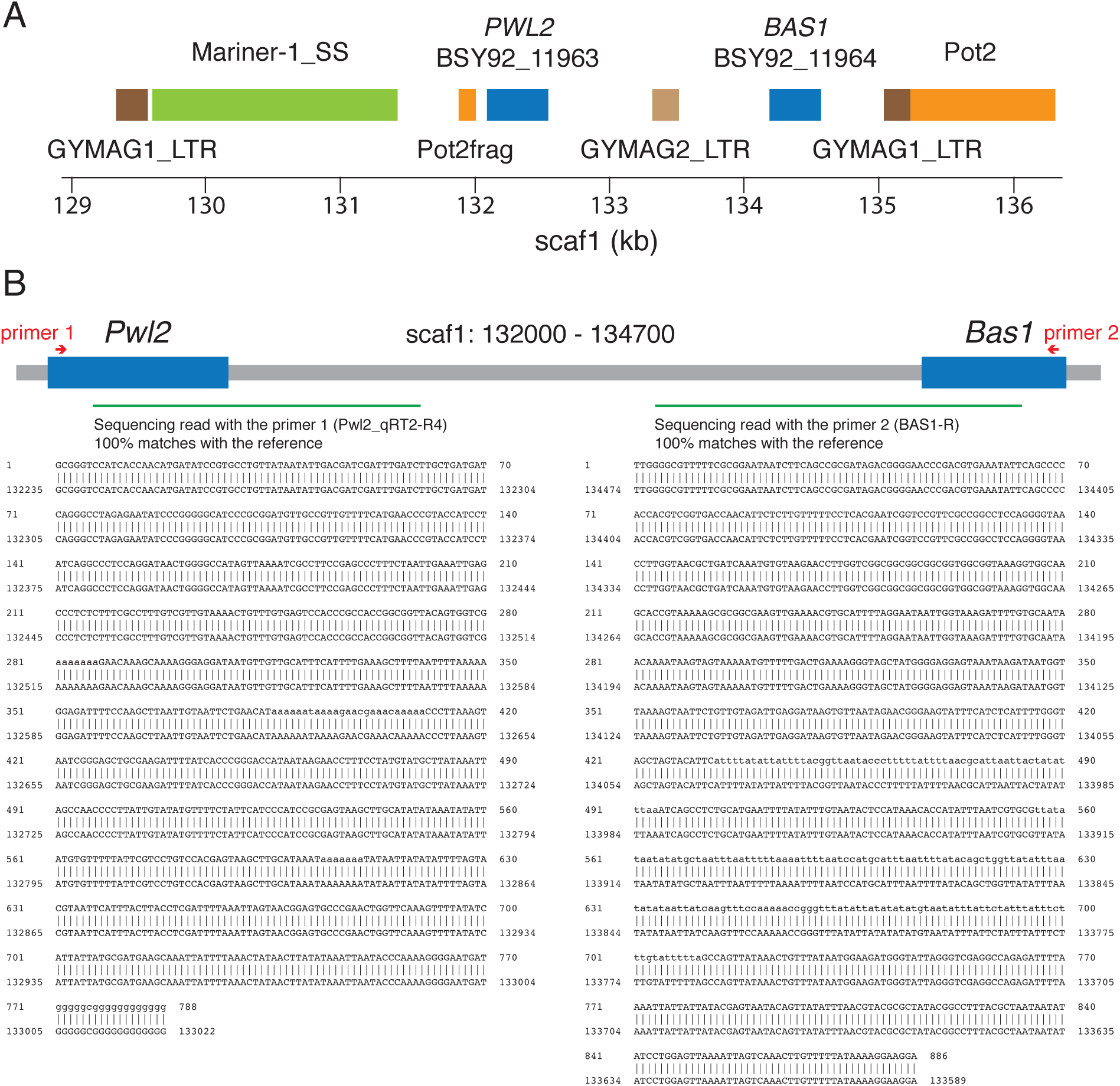
Rice blast effectors *PWL2* and *BAS1*, which are on different chromosomes in MoO strains, are side-by-side on the B71 mini-chromosome. **(A)** Location of *PWL2* and *BAS1* with flanking transposons labeled. Pot2frag is a partial Pot2 sequence. Additional transposon sequences are from MAGGY, a Gypsy-like LTR retrotransposon (GYMAG1_LTR) and a DNA transposon (Mariner-1_SS) from the TcMar-Fot1 subclass. **(B)** Validation of the neighboring structure of *PWL2* and *BAS1* via Sanger sequencing. The PCR product using the primers Pwl2_qRT2-R4 (primer 1) and BAS1-R (primer 2) was sequenced using these two primers separately. Green lines indicate the alignment regions on the scaf1 for two sequencing reads. Detailed alignments of two sequencing reads were shown underneath each green line.

**Fig. S5.**
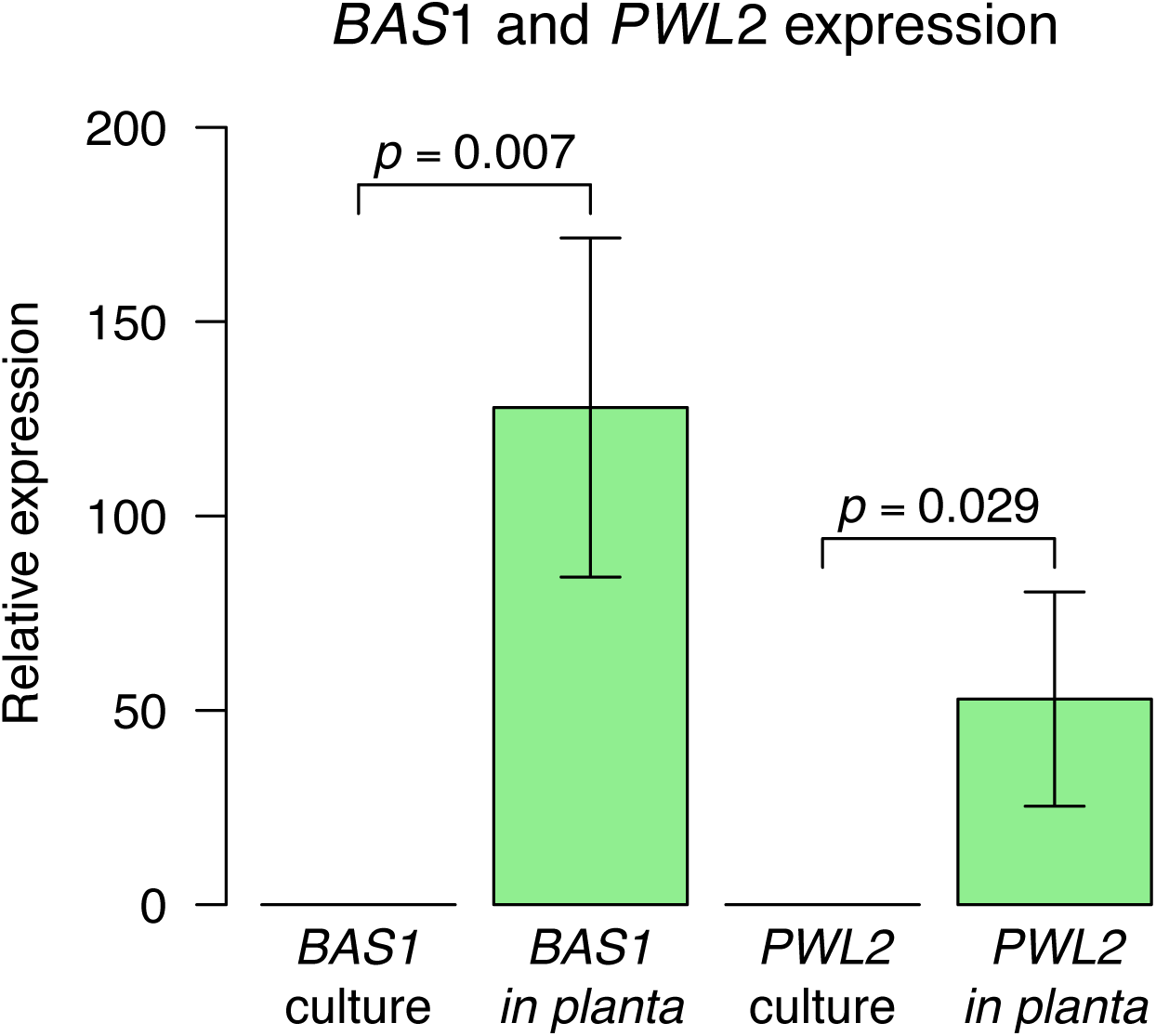
*in planta*-specific expression of *PWL2* and *BAS1* on the B71 mini-chromosome. RNA was extracted from B71-inoculated wheat leaves at 36 h after inoculation on 9-day wheat seedlings and from B71 grown in axenic culture. Expression was measured using qRT-PCR (**Methods**). Three biological replicates were used. Standard deviation is shown on each bar. P-values on top of each pair, cultured samples versus *in planta* samples, are from t-tests of expression between the two groups.

**Fig. S6.**
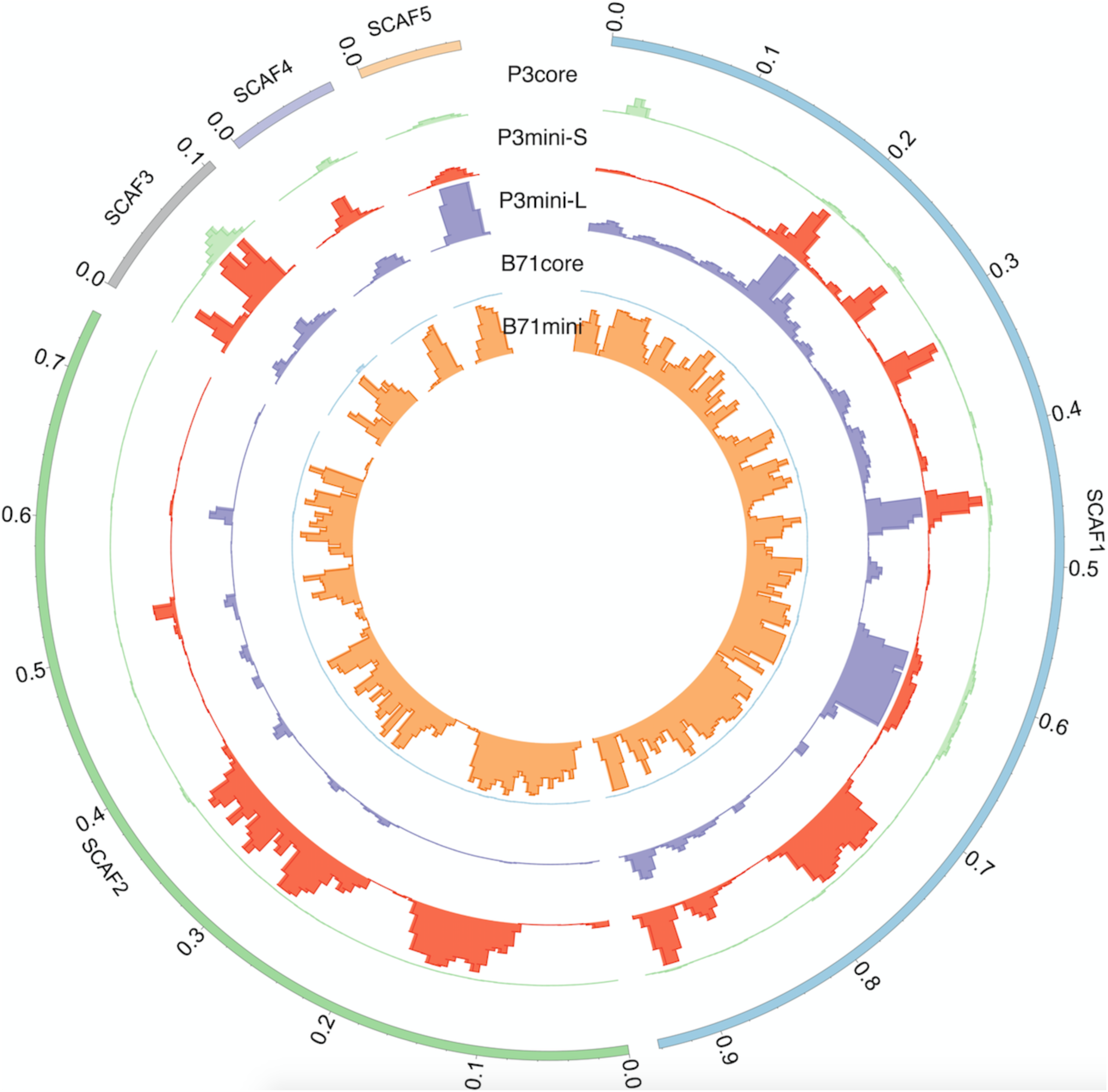
Read depths of core and mini-chromosomes on the five B71 scaffolds highlight differences among the B71 and P3 mini-chromosomes. Distributions of read depths of 10kb bins from WGS sequencing of gel-excised core or mini-chromosomes on the B71 mini-chromosome (scaf1-5). Only uniquely mapped reads were used to determine read depths. For each track, the 99.75% percentile of read depths per bin from the whole genome was used to set track height. Note the track heights are different from those of **Fig. 4D**. The change is to emphasize the high variation of read depths on the P3 large mini-chromosome.

**Fig. S7.**
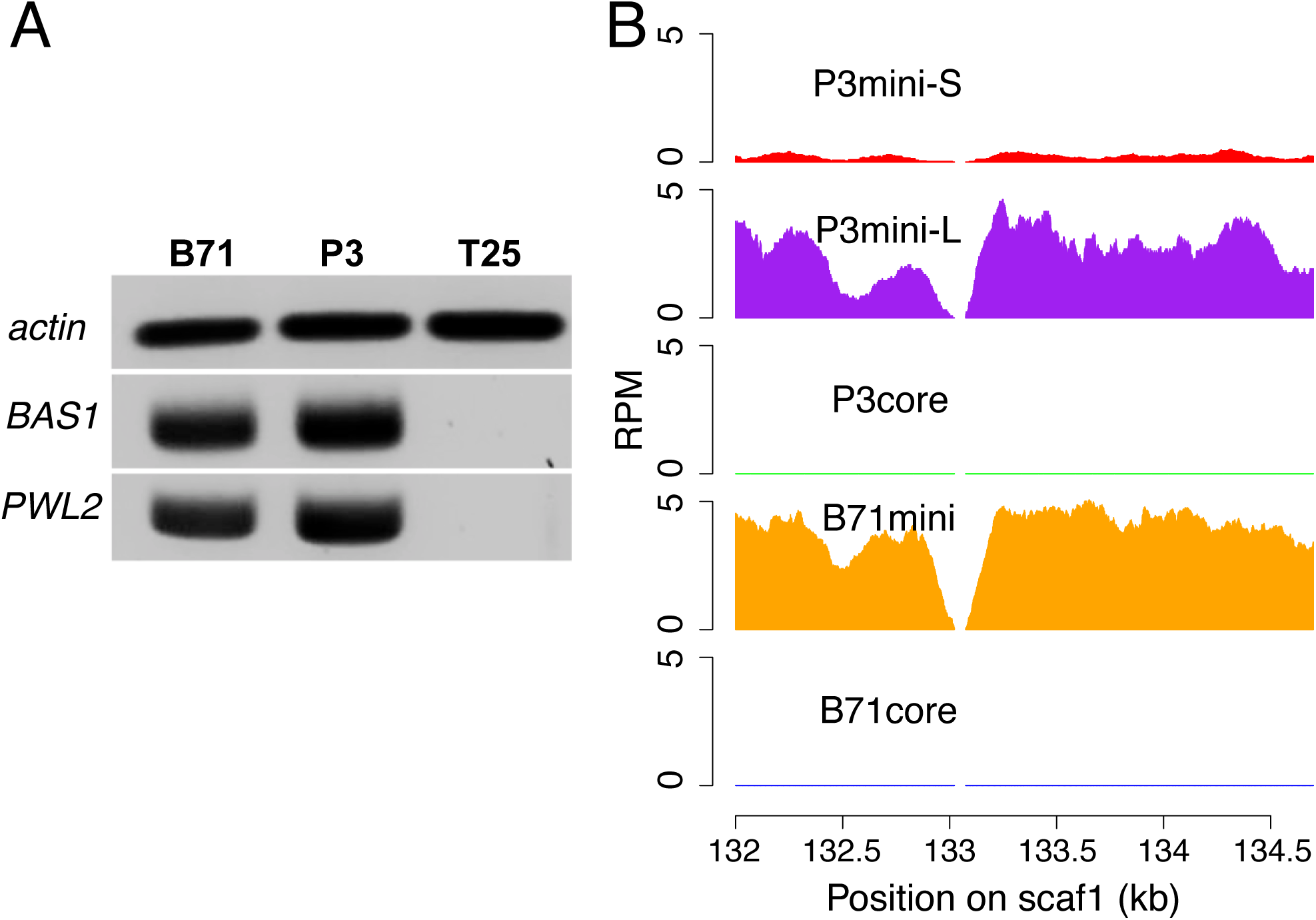
*BAS1* and *PWL2* are found on the B71 and P3L mini-chromosomes and not on the core chromosomes of B71, P3 and T25. **(A)** Genomic PCR amplification of *BAS1* and *PWL2*. Genomic DNAs of strains B71, P3 and T25 were used as templates to perform PCR (25 cycles) with primers of each of *actin*, *BAS1*, and *PWL2* genes. **(B)** Read depths of CHEF DNA sequencing on *PWL2* -*BAS1*. Distributions of read depths from WGS sequencing of gel-excised core or mini-chromosomes in the *PWL2* - *BAS1* region. Only uniquely mapped reads were used to determine read depths. RPM represents reads per million of total reads. Due to different sizes of core and mini-chromosomes, each RPM was normalized by multiplying the ratio of the total chromosome size (e.g., the size of B71mini) to the B71 core genome size. The total chromosome sizes of B71core, B71mini, P3core, P3mini-L, and P3mini-S are 43 Mb, 2 Mb, 43 Mb, 3 Mb, and 1.5 Mb, respectively.

**Fig. S8.**
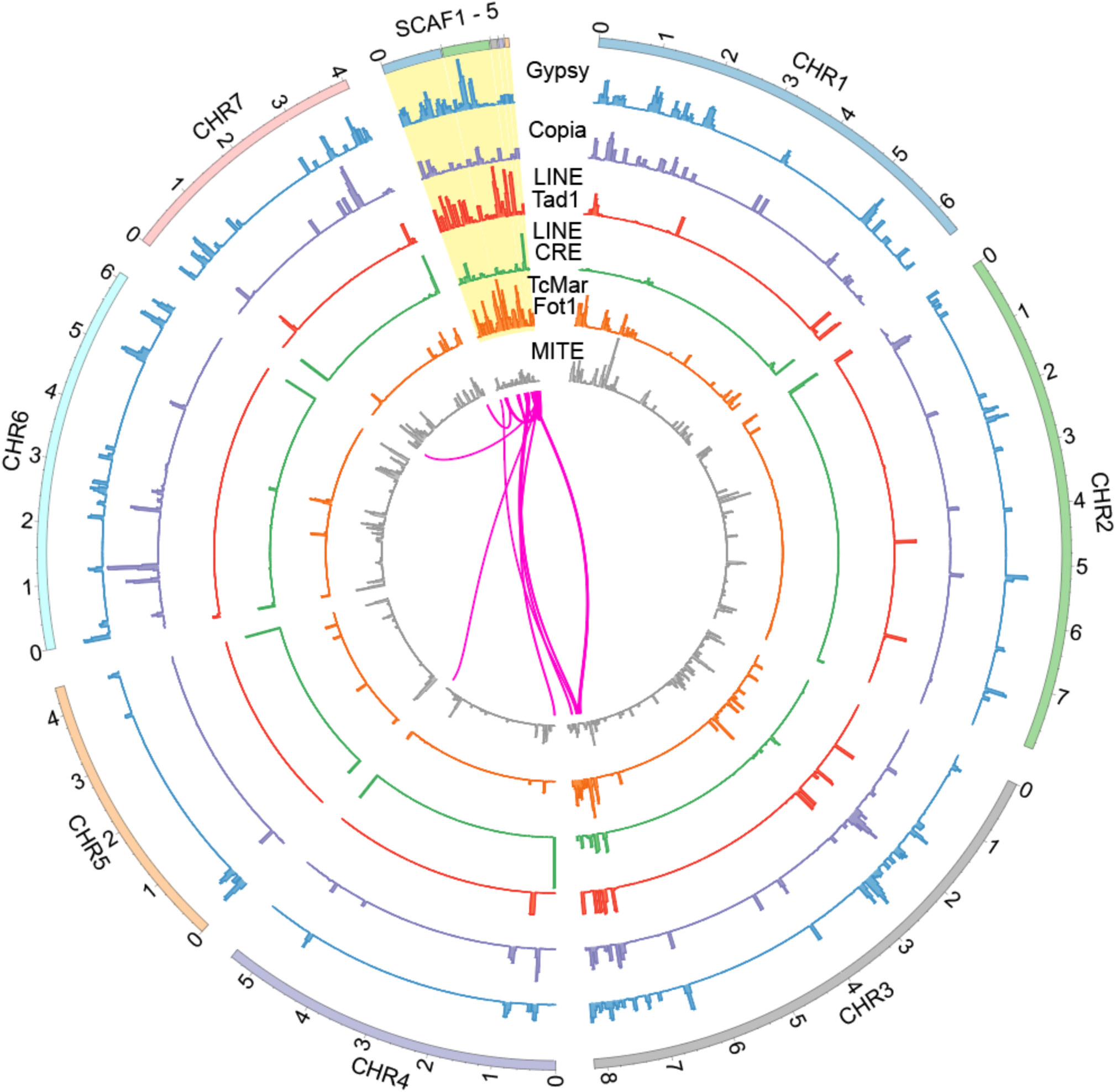
Distribution of selected transposon subclasses across the genome. The inset in the center shows large genome duplications (>10 kb and >95% identity) within five scaffolds, as well as between the scaffolds and the chromosomes 1-7. LINE subclass Tad1 contains the previously characterized retrotransposon MGR583 and subclass CRE contains the telomere-targeted retrotransposon MoTeR. The DNA transposon subclass TcMar-Fot1 contains previously characterized Pot2.

**Fig. S9.**
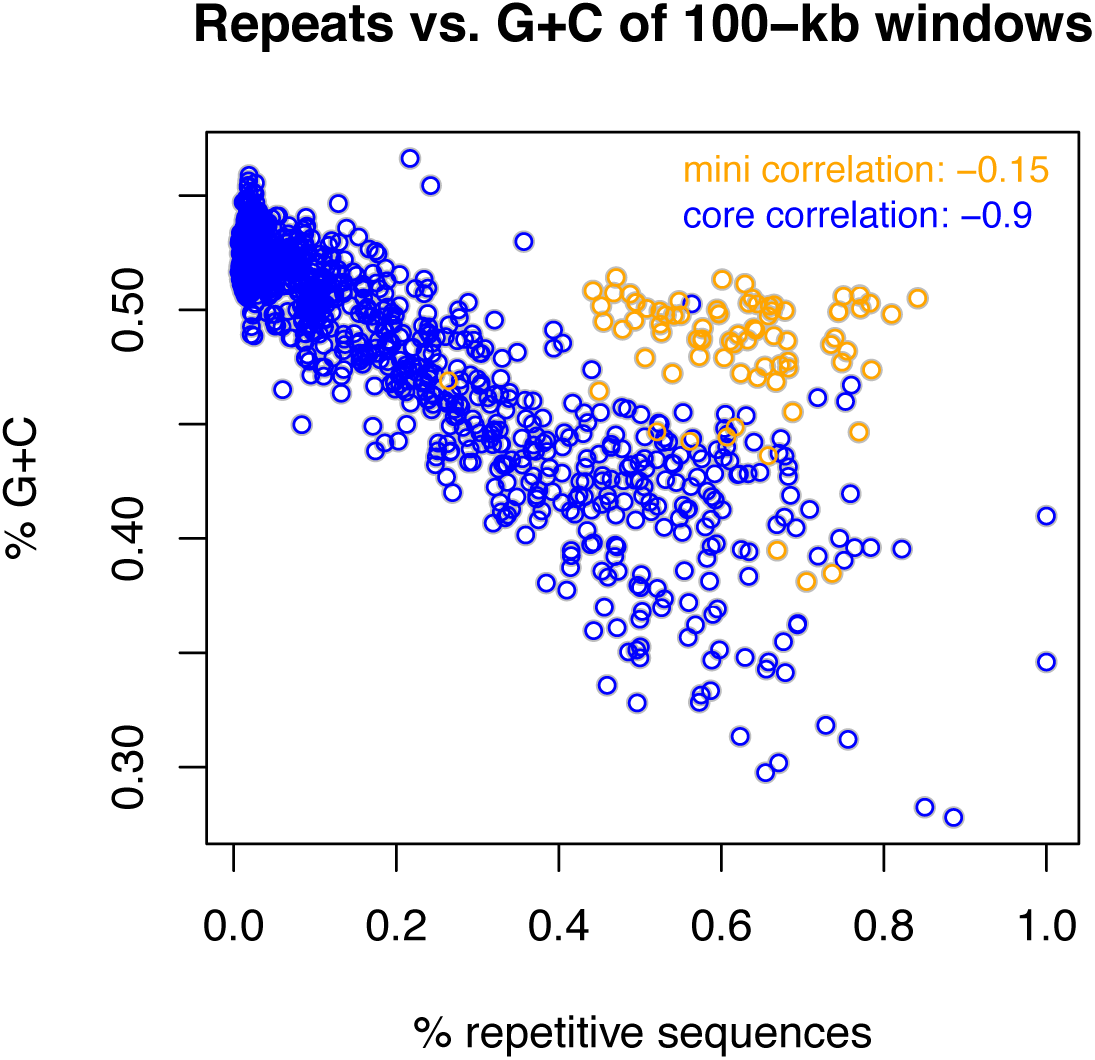
Scatter plot of GC percentages versus proportions of repetitive sequences of 100 kb bins. Orange and blue circles represent 100-kb bins from the B71 mini-chromosome (five scaffolds) and core chromosomes, respectively. Pearson correlations between GC percentages and proportions of repetitive sequences of 100 kb non-overlap genomic bins of the mini-chromosome and core chromosomes.

**Fig. S10.**
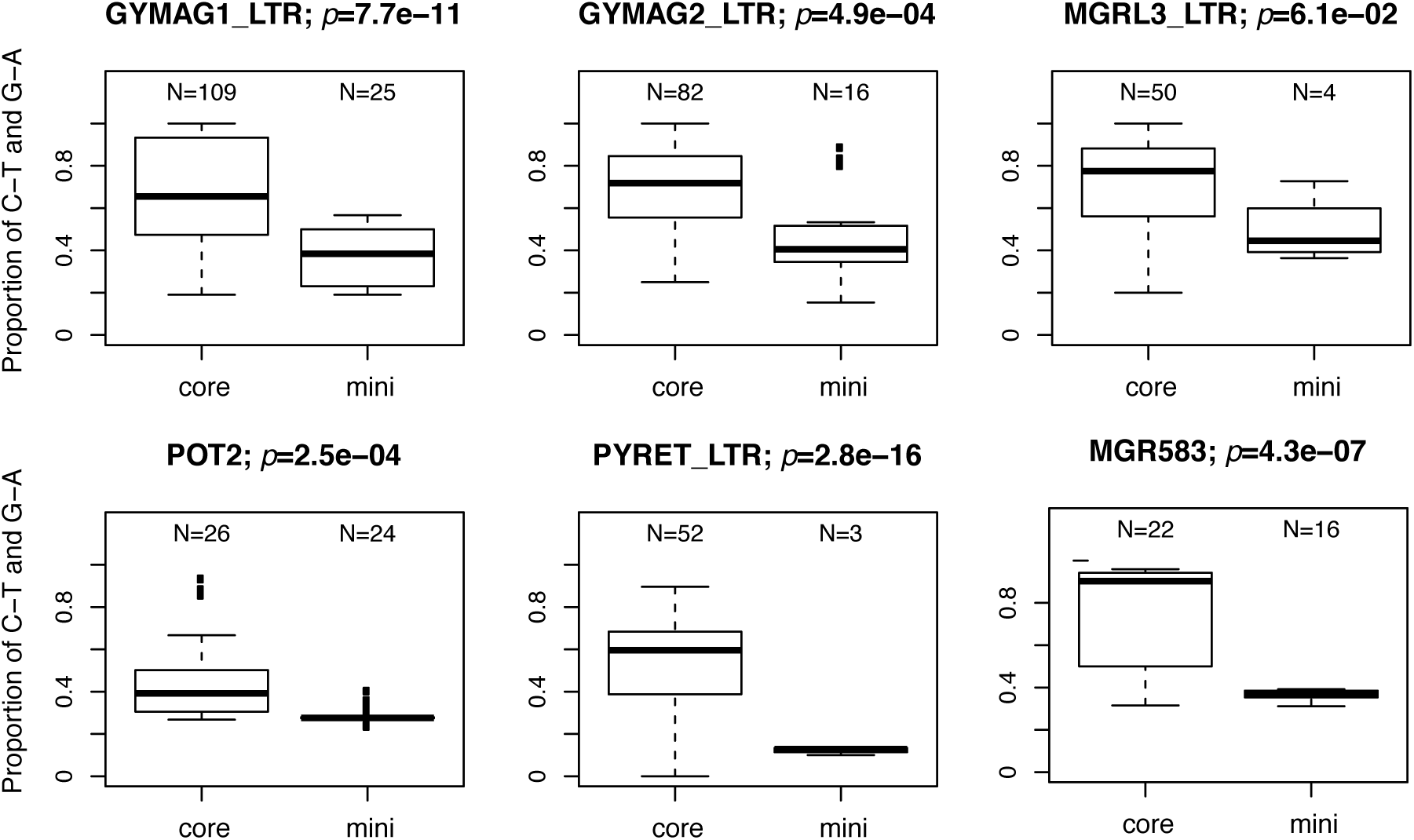
Boxplots of proportions of RIP-type changes in multiple transposable elements. Genomic sequences of each transposable element were aligned to corresponding transposon sequences from the RepeatMasker database as the reference sequences. Polymorphisms were determined for each sequence that exhibits at least 60% overlap with the reference sequence. For each transposon element, a t-test was performed to test the null hypothesis that the mean proportions of RIP-type variants out of the total mismatches of transposons located in core chromosomes was not different from that of transposons located at the mini-chromosome. P-values of t-tests were shown on the top of each boxplot.

**Fig. S11.**
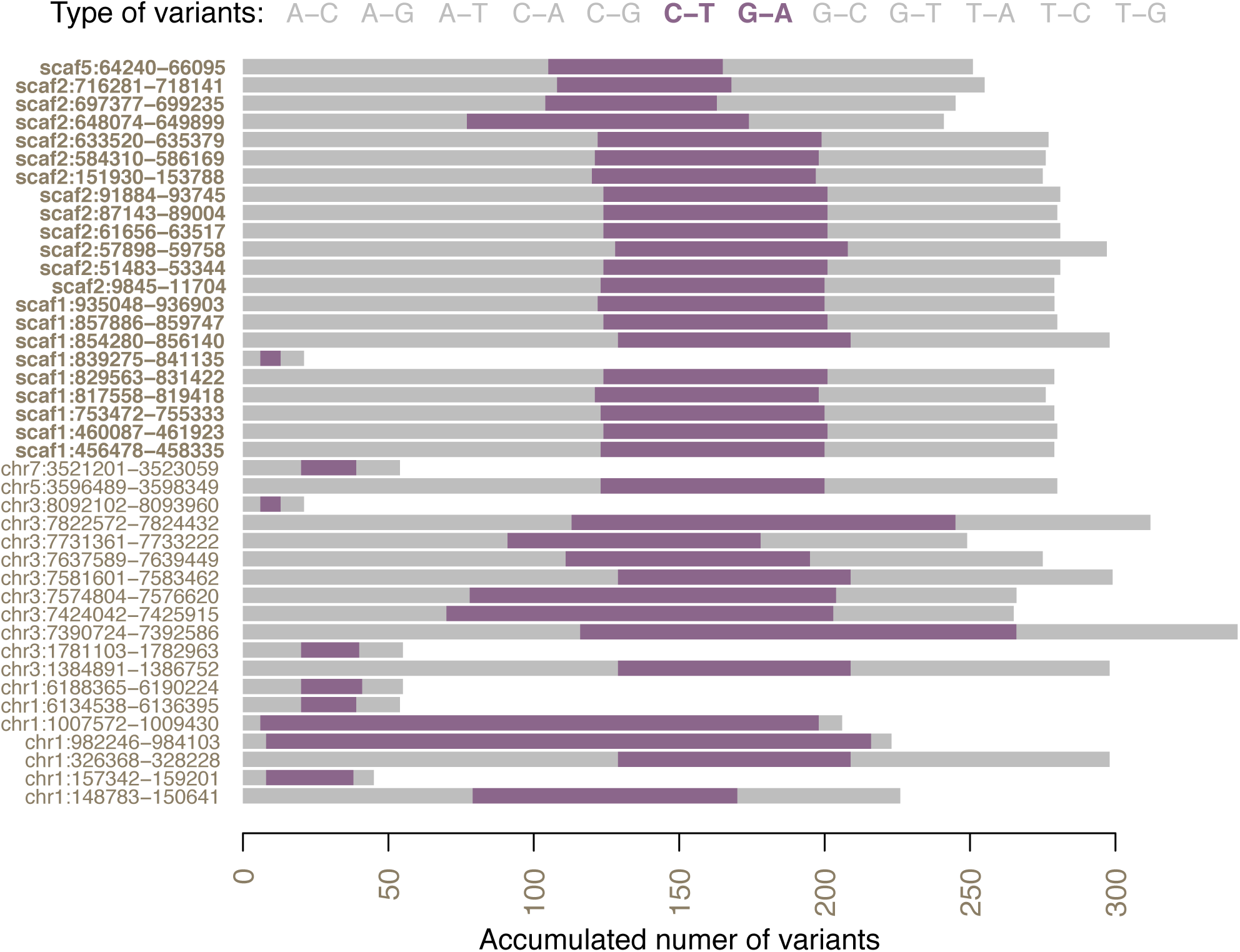
Distribution of different variant types on Pot2 homologs. Sequences of Pot2 homologs were aligned with the reference Pot2. Mismatching variants of each Pot2 homolog relative to the reference Pot2 were categorized based on nucleotide changes. All twelve variant types were listed on the top. For example, A-C represents base A on the reference Pot2 is changed to base C on Pot2 homologs. Each row shows the accumulated number of variants of a Pot2 homolog at the order of type of variation listed on the top. Two RIP-type mutations were highlighted in purple. Labels on the left show genomic locations of each Pot2 homolog.

**Fig. S12.**
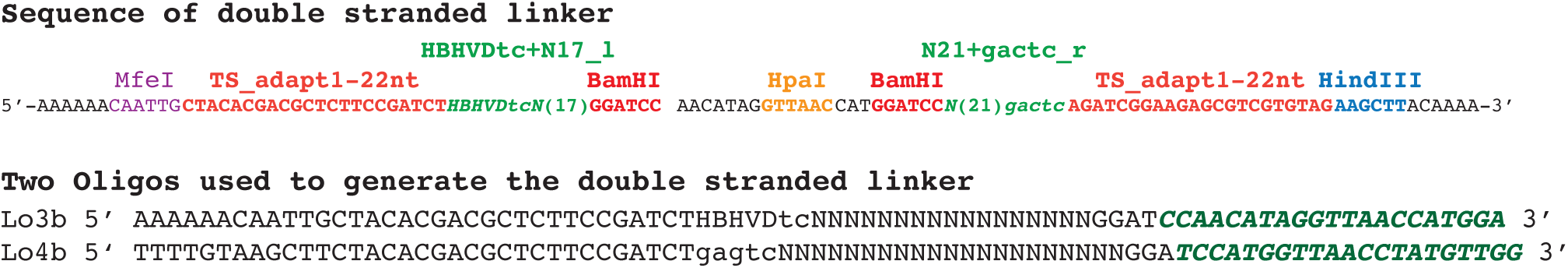
Linker design for LIEP. Two synthetic oligos with random barcodes and Illumina compatible sequence were annealed by 21 bp overlapping sequence (green italic sequences of Lo3b and Lo4b). The annealed product was then filled to form a double-stranded linker DNA (top sequence). The design of the link was shown. N(17) and N(21) indicated 17 and 21 randomly synthesized nucleotides, respectively. The linker sequence contains other IUPAC nucleotide code (e.g., H=A, C or T).

**Table Sl.**
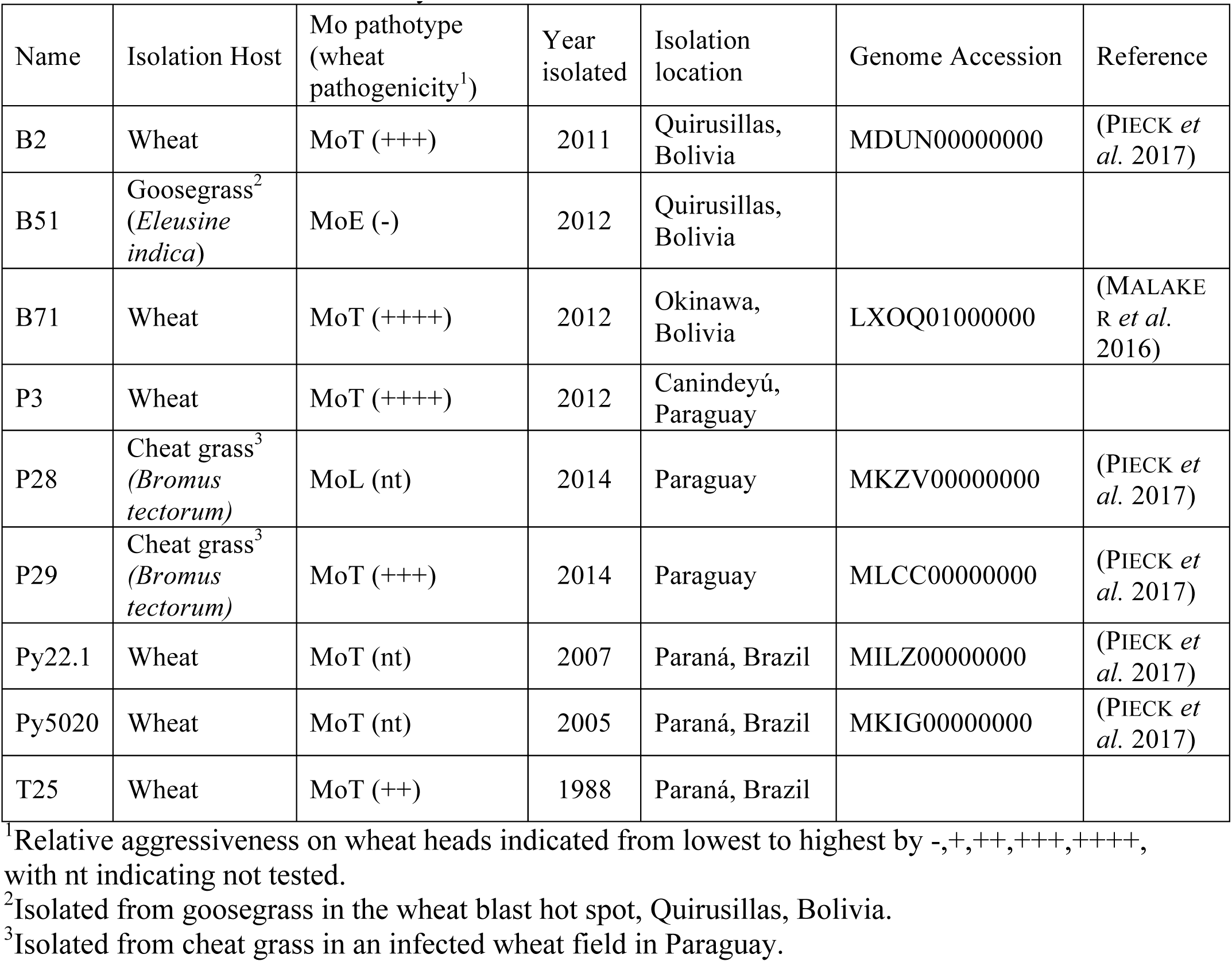
Strains used in this study

**Table S2.** Statistics of PacBio and Illumina assemblies

| **Item** | **PacBio** | **Illumina** |
| --- | --- | --- |
| Minimum cutoff of contig size (bp) | - | 500 |
| **total contig number** | **31** | **2,924** |
| total contig length (bp) | 44,480,730 | 43,297,828 |
| mean contig length (bp) | 1,434,862 | 14,807 |
| median contig length (bp) | 78,719 | 7,460 |
| SD of contig length (bp) | 436,064 | 369 |
| longest contig (bp) | 7,895,253 | 179,619 |
| shortest contig (bp) | 12,906 | 500 |
| **N50 (bp)** | **5,396,842** | **31,133** |
| L50 | 4 | 394 |
| Contigs>=1kbp | 31 | 2,671 |
| Contigs>=5kbp | 31 | 1,720 |
| Contigs>=10kbp | 31 | 1,234 |
| Contigs>=50kbp | 21 | 166 |
| **overall GC** | **0.498** | **0.499** |
| mean contig GC | 0.471 | 0.496 |
| median contig GC | 0.491 | 0.51 |
| SD of contig GC | 0.018 | 0.001 |
| max contig GC | 0.525 | 0.644 |
| min contig GC | 0.284 | 0.205 |

**Table S3.** CNV overlapping effectors genes

| Order | Effector genes | Chr | Position | B2 | B51 | P28 | P29 | P3 | Py22.1 | Py5020 | T25 |
| --- | --- | --- | --- | --- | --- | --- | --- | --- | --- | --- | --- |
| 1 | BAS1<sup>*</sup> | chr1 | 1,228,470 | % | % | % | % | % | % | % | % |
| 2 | AVR1-CO39<sup>#</sup> | chr2 | 7,253,033 | % | % | % | % | % | % | % | % |
| 3 | AVR-Pik | chr3 | 1,324,102 | % | % | % | % | % | % | % | % |
| 4 | PWL4 | chr3 | 388,420 | % | % | % | % | % | % | % | % |
| 5 | AVR-Pii | chr3 | 7,735,856 | NA | CNminus | & | % | NA | % | NA | % |
| 6 | AVR-Pi54 | chr4 | 1,079,049 | % | % | % | % | % | % | % | % |
| 7 | BAS2 | chr4 | 5,359,621 | % | % | % | % | % | % | % | % |
| 8 | BAS4 | chr5 | 17,696 | % | % | % | % | % | % | % | % |
| 9 | BAS3 | chr5 | 3,450,049 | % | % | % | % | % | % | % | % |
| 10 | PWT3<sup>#</sup> | chr5 | 3,632,010 | NA | CNplus | % | % | % | % | % | % |
| 11 | AVR-Pib | chr6 | 6,027,571 | CNplus | CNminus | % | CNplus | CNplus | CNplus | % | % |
| 12 | AVR-Pi9 | chr7 | 2,736,023 | % | % | % | % | % | % | % | % |
| 13 | AVRPiz-t | chr7 | 3,041,628 | % | % | % | % | % | % | % | % |
| 14 | AVR-Pita3 | chr7 | 344,264 | %<sup>*</sup> | % | NA | CNminus | %<sup>*</sup> | % | % | %<sup>*</sup> |
| 15 | AVR-Pik_km_kp | chr7 | 3,846,519 | % | % | % | % | CNplus | % | % | % |
| 16 | PWL2 | scaf1 | 132,123 | & | & | % | CNminus | % | & | % | CNminus |
| 17 | BAS1<sup>**</sup> | scaf1 | 134,228 | & | & | % | CNminus | % | & | % | CNminus |
| 18 | AVR-Pii | scaf1 | 29,460 | & | NA | NA | & | % | & | CNplus | & |

%. CNequal, conserved between the strain and B71
&: The CNV index was between CNequal and CNminus and was referred to as "polymorphic"
%<sup>\*</sup>: CNequal but partial genes were in polymorphic regions
NA: no CNV segments overlapping with gene regions of effectors
\*: 68% identity to 70-15 BAS1
\*\* : 99% identity to 70-15 BAS1

**Table S4.**
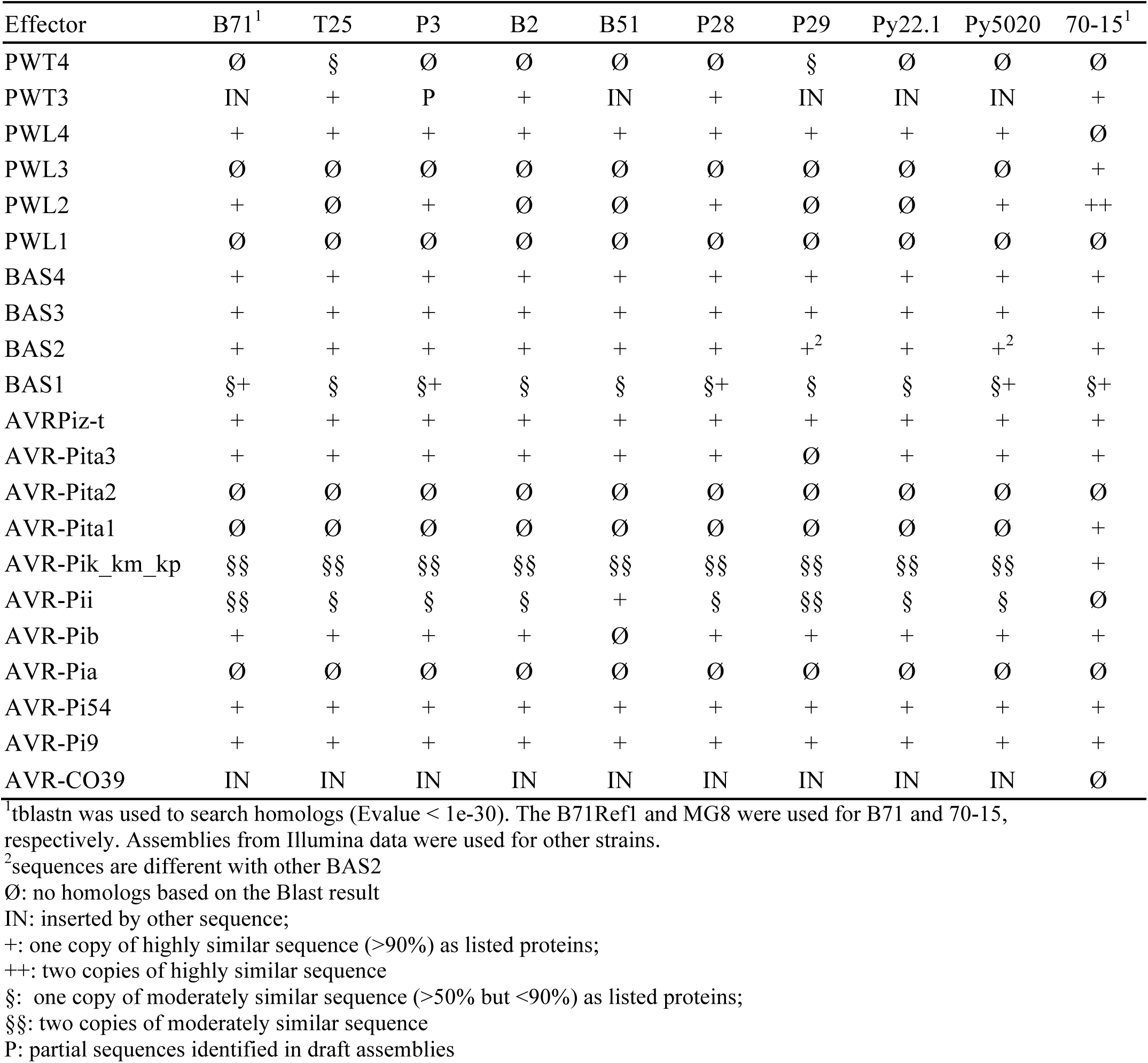
Copy number of effectors based on assembled sequences

**Table S5.** Summary of repetitive elements of the B71 MoT genome

| Repeats | Counts | Total (bp) / (%<br>of genome) | Counts<br>in core | Total (bp) in core<br>/ (% of core) | Counts<br>in mini | Total in mini (bp)<br>/ (% of mini) |
| --- | --- | --- | --- | --- | --- | --- |
| Class I (LTR) |  |  |  |  |  |  |
| Copia | 698 | 554,832/1.25 | 594 | 496,082/1.17 | 104 | 58,750/3.04 |
| Gypsy | 1,322 | 1,678,110/3.77 | 1,040 | 1,411,961/3.32 | 282 | 266,149/13.79 |
| unknown | 692 | 626,450/1.41 | 590 | 537,936/1.26 | 102 | 88,514/4.59 |
| Class I (LINE) |  |  |  |  |  |  |
| Tad1 | 165 | 249,837/0.56 | 94 | 141,286/0.33 | 71 | 108,551/5.62 |
| CRE | 143 | 172,313/0.39 | 101 | 138,429/0.33 | 42 | 33,884/1.76 |
| Jockey | 85 | 65,307/0.15 | 55 | 35,249/0.08 | 30 | 30,058/1.56 |
| I | 92 | 22,508/0.05 | 59 | 8,882/0.02 | 33 | 13,626/0.71 |
| R2 | 26 | 14,163/0.03 | 22 | 13,661/0.03 | 4 | 502/0.03 |
| Class I (SINE) |  |  |  |  |  |  |
| Alu | 192 | 71,583/0.16 | 148 | 37,856/0.09 | 44 | 33,727/1.75 |
| Mg | 1 | 195/0.0004 | 0 | 0/0 | 1 | 195/0.0101 |
| Class II (DNA) |  |  |  |  |  |  |
| TcMar-Fot1 | 395 | 306,805/0.69 | 282 | 197,036/0.46 | 113 | 109,769/5.69 |
| TcMar-Pogo | 8 | 3,135/0.01 | 0 | 0,000/0.00 | 8 | 3,135/0.16 |
| MITE | 572 | 152,506/0.34 | 528 | 144,451/0.34 | 44 | 8,055/0.42 |
| Unclassified elements | 2,209 | 1,222,299/2.75 | 1,839 | 957,562/2.25 | 370 | 264,737/13.72 |
| **<u>Total transposon</u>**<br>**<u>elements</u>** | 6,600 | 5,140,043/11.56 | 5,352 | 4,120,391/9.69 | 1,248 | 1,019,652/52.83 |
| rRNA | 25 | 46,010/0.10 | 25 | 46,010/0.11 | 0 | 0/0 |
| Satellite | 77 | 8,595/0.02 | 77 | 8,595/0.02 | 0 | 0/0 |
| Simple_repeat | 12,715 | 466,976/1.05 | 12,606 | 462,423/1.09 | 109 | 4,553/0.24 |
| Low_complexity | 1,667 | 73,416/0.17 | 1,654 | 72,867/0.17 | 13 | 549/0.03 |
| **<u>Total</u>** | 21,084 | 5,735,040/12.90 | 19,714 | 4,710,286/11.08 | 1,370 | 1,024,754/53.10 |

**Table S6.**
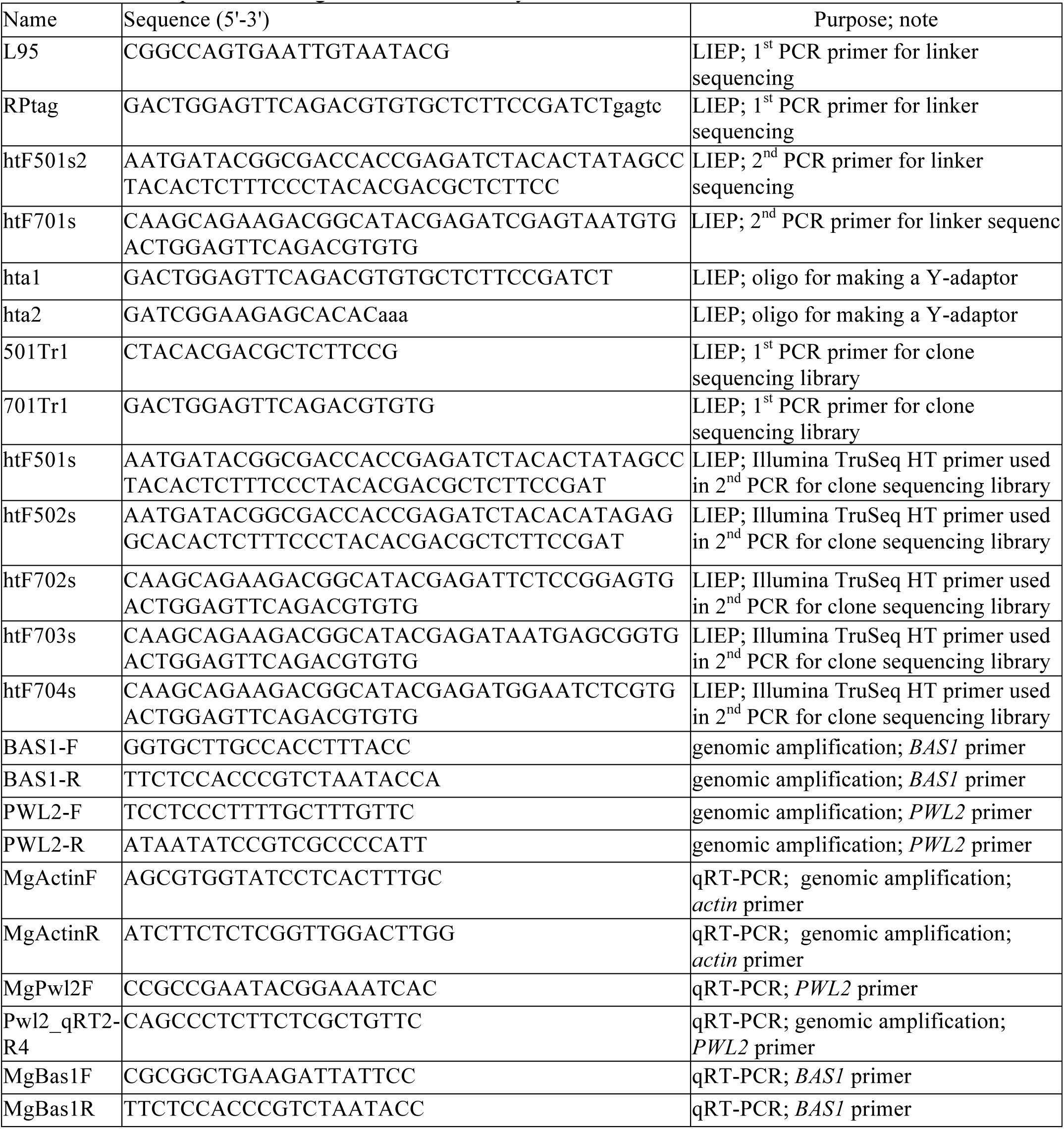
List of primers or oligos used in this study

**Data S1.** Difference between PacBio drafted sequences before Illumina correction and the B71Ref1 sequences

**Data S2.** Functional annotation of genes

**Data S3.** GFF file of genome annotation

**Data S4.** Gene expression from RNA-Seq

**Data S5.** List of putative effectors

**Data S6.** Gene ontology of genes

